# Heat shock alters the distribution and interaction of major nuclear proteins—Lamin B with DNA topoisomerase II and chromatin

**DOI:** 10.1101/2024.02.28.582469

**Authors:** Marta Rowińska, Aleksandra Tomczak, Jadwiga Jabłońska, Aleksandra Zielińska, Katarzyna Piekarowicz, Magdalena Machowska, Ryszard Rzepecki

## Abstract

Lamins and topoisomerase II (Top2) contribute to nuclear architecture and chromatin organization and gene expression, but their responses to thermal stress remain incompletely understood.

Here, we investigated heat shock-induced changes in the phosphorylation, solubility, subnuclear distribution and molecular associations of B-type lamin Dm and Top2 in Drosophila melanogaster Kc and S2 cells, embryos and third-instar larvae, including their reversibility during recovery. Heat shock induced a transient increase in lamin Dm phosphorylation at serine 25 (S25) and altered the solubility of lamin Dm, Top2 and several nuclear regulatory proteins, with particularly pronounced changes in S2 cells. Immunofluorescence analyses revealed distinct changes in Top2 distribution depending on the experimental model: the formation of intranuclear Top2-positive granules in Kc and S2 cells and increased peripheral localization of Top2 in embryonic nuclei. Chromatin redistribution toward the nuclear periphery was observed in third-instar larval nuclei. Co-immunoprecipitation experiments demonstrated an association between lamin Dm and Top2 but no increase in their direct interaction was detected following heat shock. In vivo photocrosslinking and immunoprecipitation analyses indicated enhanced association of lamin Dm and Top2 with nucleic acids under heat shock conditions, involving limited fractions of the respective protein populations. The observed heat shock-associated changes were largely reversible during recovery. Together, these findings demonstrate that thermal stress induces reversible, model-dependent changes in the biochemical properties, nuclear distribution and nucleic acid associations of lamin Dm and Top2. These observations suggest a potential contribution of both proteins to the reorganization of nuclear architecture during the heat shock response, with potential contribution to the regulation of gene expression associated with heat shock.

## 1. Introduction

Heat shock (HS) rapidly alters chromatin organization and the biochemical properties of nuclear proteins. Nuclear lamins and DNA topoisomerase II (Top2) are of particular interest because they contribute to nuclear architecture and chromatin-associated processes. Lamins form the nuclear lamina, support nuclear mechanical stability and mechanotransduction, and participate in chromatin organization and positioning, gene regulation and nucleocytoplasmic signaling [1, 2, 3]. Top2 resolves DNA topological constraints associated with transcription, replication and chromosome segregation [4, 5]. How the localization and nucleic acid associations of these proteins change during HS and recovery remains incompletely understood.

*Drosophila melanogaster* provides a useful system for addressing these questions because it has one B-type lamin gene (lamin Dm), one A-type lamin gene (lamin C) and one Top2 gene [6, 7]. Lamin Dm is expressed throughout development, whereas lamin C is developmentally regulated and appears later in embryogenesis [8]. Kc cells express both lamins, while S2 cells predominantly express lamin Dm at the protein level, allowing the responses to HS to be compared in distinct cellular backgrounds [9].

An important feature of Drosophila lamin Dm is the existence of several electrophoretically distinguishable forms generated by post-translational modification rather than by alternative splicing of the coding gene. Two developmentally regulated transcripts encode the common precursor lamin Dm0, which is rapidly converted to the phosphorylated Dm1 form; additional phosphorylation gives rise to Dm2. Dm1 and Dm2 are the main interphase, relatively insoluble forms, and their proportions vary with cellular conditions. More soluble mitotic and meiotic forms, designated DmMit and DmMei, are associated with changes in phosphorylation during nuclear lamina disassembly. Early studies from the laboratory of Paul A. Fisher established the biosynthesis and interconversion of these lamin forms and showed that HS promotes the conversion of Dm2 to Dm1 [10, 11, 12, 13].

Lamin Dm phosphorylation is heterogeneous and site specific. Residues identified as phosphorylation sites in vivo include S19, S25, S45, S595 and T597, while the C-terminal Cdk1-regulated site T435 is important for mitotic disassembly and intracellular localization. Although several sites can be modified, metabolic labeling studies suggested that the average number of phosphate groups per lamin molecule is limited [12, 14, 15, 13, 3]. Mutational studies linked phosphorylation at S45 and T435, and to a lesser extent S25 and S595, to changes in lamin Dm solubility and nuclear distribution [16]. Of particular relevance here, S25 lies within the R22PPS motif proposed to mediate binding of 14-3-3 proteins; a regulatory role for this motif remains a hypothesis rather than an established mechanism [13, 3]. These findings motivate examining S25 phosphorylation together with lamin Dm solubility and localization under HS.

Lamin Dm also has experimentally demonstrated interactions with nucleic acids. In vivo UV photocrosslinking followed by immunoprecipitation revealed associations of interphase lamin Dm with both DNA and RNA, whereas such associations were not detected for mitotic lamin Dm. In the same experimental system, no comparable nucleic acid association was detected for lamin C, emphasizing a functional distinction between the two Drosophila lamins under the conditions tested [17]. Phosphomimetic analyses further indicated that individual lamin Dm phosphorylation sites can influence its distribution and association with chromatin [16]. Thus, changes in lamin Dm phosphorylation during HS may be accompanied by changes in its nucleic acid associations, a possibility that requires direct experimental examination.

Top2 also exists in heterogeneous biochemical populations, as shown in studies by Fisher and colleagues and in subsequent work on Drosophila Kc cells and embryos [18, 19, 20, 21, 22]. Cell fractionation identified a predominantly low-speed-pellet-associated population (P-10), including chromatin-associated Top2, and a biochemically distinct population recovered from the high-speed pellet (P-130), including non-chromosomal mitotic Top2. These forms differ in solubility, phosphorylation, enzymatic activity and interactions with nucleic acids. P-130 Top2 was more extensively phosphorylated and less enzymatically active than P-10 Top2, and phosphatase treatment increased its activity. In vivo photocrosslinking showed that Top2 associates with both DNA and RNA during interphase and mitosis; the P-130 population was associated predominantly with RNA. The C-terminal region of Top2, including approximately its last 200 amino acids, was implicated in these nucleic acid interactions and is a major site of phosphorylation [22]. These findings indicate that Top2 cannot be regarded as a single homogeneous nuclear population.

Importantly, earlier in vivo studies demonstrated that HS changes the association of Top2 with specific chromatin regions. Mapping of Top2-mediated DNA cleavage sites in Drosophila heat-shock chromatin revealed substantial changes in the intensity of selected cleavage signals upon induction [23]. Analysis of the 87A7 locus further showed a pronounced redistribution of Top2 sites of action when hsp70 genes were activated, followed by another redistribution during recovery as chromatin recondensed [24]. Thus, HS can enhance Top2-associated signals at particular sites and reorganize its chromatin interactions, although these locus-specific findings do not establish a genome-wide increase in Top2 binding. Earlier extraction experiments also reported HS-dependent changes in the retention of nuclear proteins in operationally defined karyoskeletal fractions [25, 26], but such fractions do not establish the precise localization or direct interactions of these proteins in intact nuclei.

Together, these studies suggest that the HS response may involve coordinated changes in the biochemical states, nuclear distributions, and nucleic acid associations of lamin Dm and Top2. Here, we examined lamin Dm S25 phosphorylation, protein extractability, Top2 localization, lamin Dm–Top2 association and interactions with nucleic acids during HS and recovery in Drosophila Kc and S2 cells, embryos and third-instar larvae. By comparing biochemical and imaging results across models, we sought to distinguish directly observed stress-associated changes from possible functional consequences, without presuming that protein redistribution demonstrates altered gene expression or chromatin tethering.

## 2. Materials and Methods

### Antibodies

The affinity-purified rabbit anti-lamin Dm and domain-specific rabbit anti-Top2 antibodies used in this study have been described previously [22]. Guinea pig anti-LBR antibodies were kindly provided by Georg Krohne (University of Würzburg, Germany; [27]). The mouse monoclonal antibodies ADL67.10 (anti-lamin Dm, recognizing a fragment of the lamin rod domain), ADL84.12 (anti-lamin Dm, specific for the nonphosphorylated S25 residue), and LC28.26 (anti-lamin C) were obtained from the Developmental Studies Hybridoma Bank (DSHB). Mouse and rabbit anti-α-tubulin antibodies were obtained from DSHB (clone 12G10) and Thermo Fisher (#PA5-29444), respectively. The rat monoclonal anti-Hsp70 antibody (heat shock protein 70 kDa; #NBP2-59342) and rabbit polyclonal anti-HDAC1 antibody (histone deacetylase 1; #NB500-124) were obtained from Novus Biologicals. The rabbit polyclonal anti-heat shock factor (Hsf) antibody was custom-made by Thermo Scientific. Additional immunofluorescence staining was performed using rat anti-Hsp90 (heat shock protein 90 kDa; #ab13494) and rabbit anti-Pol2PS5 (RNA polymerase II phosphorylated at Ser5; #ab5131) from Abcam, and rabbit anti-HP1 (heterochromatin protein 1; #NB110-40623) from Novus Biologicals. All secondary antibodies were obtained from Jackson ImmunoResearch. HRP-conjugated anti-mouse (#715-035-151), anti-rabbit (#111-035-144), and anti-rat (#712-035-153) antibodies were used to detect antigens on immunoblots. For immunofluorescence, anti-mouse antibodies were conjugated with TRITC or DyLight649 (#715-025-151 and #715-495-150), anti-rat antibodies with TRITC (#712-025-153), and anti-rabbit antibodies with Alexa488 (#711-545-152).

### Drosophila melanogaster embryos and cell cultures

The D. melanogaster Oregon R-P2 strain (RRID:BDSC_2376) was obtained from the Bloomington Drosophila Stock Center (Bloomington, USA; NIH P40OD018537). Embryos were collected using egg-laying cages. Flies were transferred from standard medium (6.25% [w/v] corn grits, 3.13% [w/v] corn flour, 6.25% [w/v] sugar, 12.5% [w/v] yeast, and 0.5% [w/v] bacteriological agar) to cages containing embryo-collection medium (20 mL apple juice, 10 mL cherry syrup, 10 mL water, 2.5 g sugar, and 1.6 g bacteriological agar) coated with yeast paste. Flies were allowed to lay eggs for 11 h, after which a 1-h heat shock at 37 °C was applied. The analyzed embryos were 1–12 h old. Kc cells (ECACC #90070550) and S2 cells (Drosophila Genomics Resource Center, stock 6; https://dgrc.bio.indiana.edu//stock/6; RRID:CVCL_TZ72; kindly provided by the Department of Cytobiochemistry, University of Wroclaw, Poland) were cultured in Schneider’s Drosophila Medium (Gibco #21720024) supplemented with 10% fetal bovine serum (Sigma #7524) and 1% antibiotic–antimycotic mixture (Gibco #15240062) at 23 °C under normal conditions (N), without supplemental CO.

### Preparation of D. melanogaster embryos and cultured cells for Western blotting

Embryos were transferred to strainers fitted with NITEX nylon mesh (40 µm) and rinsed with water. The chorion was removed using 5% sodium hypochlorite. Embryos were then washed with buffer containing 5 mM NaCl, 20 mM Tris-Cl (pH 7.5), and freshly added protease and phosphatase inhibitors. They were subsequently processed according to the protocols for the respective experiments.

Cells were seeded in 10-cm culture dishes 24 h before each experiment. They were collected by gentle scraping and centrifugation. Cell pellets were resuspended in Laemmli sample buffer and heated at 95 °C for 10 min before Western blot analysis. To induce heat shock (HS), embryos, larvae, or cultured cells were transferred to 37 °C for 1 h, unless a different duration is specified. Recovery (R) was initiated by returning samples to 23 °C for an additional 6, 24, 48, 72, or 96 h, as indicated. Unless otherwise stated, embryos and larvae were exposed to 37 °C for 1 h and then allowed to recover at 23 °C for 4 h.

### Western blot analysis

Proteins were separated by SDS–PAGE on 7–12% polyacrylamide gels, depending on the experiment, according to Laemmli [28], and transferred electrophoretically to nitrocellulose membranes (0.45 µm; Amersham Protran) using a Bio-Rad Mini Trans-Blot Cell, as described by Harlow and Lane [29]. Protein molecular weights were estimated using a prestained protein ladder (Thermo Fisher #26616). After blocking for 1 h in 5% nonfat milk prepared in PBST (phosphate-buffered saline [PBS] containing 0.075% Tween-20), membranes were incubated with primary antibodies overnight at 4 °C, washed three times with PBST, and incubated with HRP-conjugated secondary antibodies for 1.5 h at room temperature (RT). Signals were detected using SuperSignal™ West Pico PLUS Chemiluminescent Substrate (Thermo Fisher #34580) or Clarity Western ECL Substrate (Bio-Rad #1705061). Images were acquired with the ChemiDoc MP Imaging System (Bio-Rad) and quantified using Image Lab software, version 5.2 (Bio-Rad).

For colorimetric detection, goat anti-IgG antibodies conjugated to calf intestinal alkaline phosphatase and/or donkey anti-IgG antibodies conjugated to horseradish peroxidase were used with the corresponding chromogenic substrates (Jackson ImmunoResearch).

In some co-immunoprecipitation (co-IP) experiments involving lamins and Top2, including experiments with BrdU and/or BrdC labeling, UV crosslinking, and radioactive ³²P labeling, both proteins were detected simultaneously using colorimetric methods. Secondary antibodies conjugated to horseradish peroxidase or alkaline phosphatase were used with suitable chromogenic substrates. The antibodies specified for the two targets were goat anti-rabbit IgG conjugated to phosphatase for Top2 and goat anti-mouse IgG conjugated to alkaline phosphatase for lamin Dm. Control IP and co-IP experiments used IgG isolated from preimmune rabbit serum or control mouse IgG.

### Immunofluorescence

Cells were grown on gelatin-coated coverslips for 24 h and, where indicated, exposed to heat shock before fixation. Cells were fixed with 4% paraformaldehyde, permeabilized with 0.5% Triton X-100 in PBS, and blocked with 5% fetal bovine serum in PBS. Cells and embryos were processed as described previously [17, 30].

Before fixation, embryos were dechorionated with 5% sodium hypochlorite for 3 min and rinsed with water. Typically, 100 µL of packed embryos were fixed in a mixture of 300 µL of 4% paraformaldehyde and 1 mL of heptane for 20 min at RT. The fixative phase was then replaced with 1 mL of methanol. Embryos were washed twice with PBS, rehydrated in PBS containing 0.1% Triton X-100 for 30 min at RT, and permeabilized in PBS containing 0.5% Triton X-100 for 30 min at RT with stirring. Finally, embryos were washed with PBS and stained as described below.

Cells and embryos were incubated with primary antibodies overnight at 4 °C and then with secondary antibodies for 1.5 h at RT. Stage 10–12 embryos are shown in Figure 3. The proportion of nuclei exhibiting peripheral Top2 localization was determined by manual counting. Nuclei with irregular Top2 staining or reduced staining in the nuclear center were counted and expressed as a percentage of all visible nuclei under normal (N) and HS conditions. Differences between groups were evaluated using the chi-square test. The same counting procedure was used to assess peripheral redistribution of chromatin, as visualized by DAPI staining.

Proximity ligation assay (PLA) staining was performed using the Duolink PLA Multicolor Reagent Pack (Sigma–Aldrich #DUO96000) according to the manufacturer’s instructions. Samples were incubated with primary antibodies for 16 h at 4 °C, washed with PBS, and incubated with secondary antibodies in PBS for 1.5 h at RT. After washing, slides were mounted in VECTASHIELD medium containing DAPI. PLA was performed on Kc cells and third-instar larval preparations; larval smears were prepared and processed in the same way as cultured cells. For quantification, the number of visible PLA-positive foci or clusters was divided by the number of nuclei identified by DAPI staining, assuming one nucleus per cell.

Confocal images were acquired using a Leica SP8 microscope and processed with Fiji image-analysis software.

### Solubility assay

#### Kc and S2 cells

Cells were seeded 24 h before the experiment at a density of 2.5 × 10 cells/mL. After heat shock, cells were detached by scraping, transferred to 15-mL tubes, collected by centrifugation at 500 × g for 5 min, and washed with PBS. The pellets were resuspended in extraction buffer containing 5 mM MgCl, 50 mM NaCl, 50 mM Tris-HCl (pH 7.5), 250 mM sucrose, 0.1 M EDTA, 1% Triton X-100, and freshly added protease and phosphatase inhibitors. After mechanical homogenization, the lysate was incubated on ice for 20 min. An aliquot of the lysate was collected as a control before further extraction. The remaining lysate was centrifuged at 10,000 × g for 10 min at 4 °C, and the supernatant (50 mM NaCl-soluble fraction) was collected. The pellet was resuspended in the same extraction buffer containing 150 mM NaCl, incubated on ice for 20 min, and centrifuged; the supernatant (150 mM NaCl-soluble fraction) was collected. The procedure was repeated with buffer containing 500 mM NaCl. The collected samples were heated in denaturing buffer (5% DTT and 40 mM DTT) at 96 °C for 10 min and prepared for SDS–PAGE in standard Laemmli sample buffer. A 10-µL aliquot of each sample was loaded onto the gel, irrespective of the fraction.

#### Embryos

Collected and washed embryos were resuspended in the same extraction buffer used for cultured cells, at a ratio of 25 mg of embryos to 125 µL of buffer. Subsequent extraction steps were performed as described above.

#### Statistics

Mean band intensities were compared between samples maintained under normal conditions (N; 23 °C) and after heat shock (HS; 1 h at 37 °C). For each fraction (lysate, 50 mM, 150 mM, and 500 mM NaCl), an independent Student’s t-test was performed. The analysis used mean values calculated from three biological replicates and at least four technical replicates.

RNA extraction and first-strand cDNA synthesis

Semi-adherent Kc cells were seeded at a density of 2.5 × 10 cells/mL in Schneider’s medium using 6-cm culture dishes (1.5 × 10 cells in total). After 24 h, the medium was removed, and cells were gently washed once with PBS. Lysis buffer from the Universal RNA Purification Kit (EURx) was added, and the remaining steps were performed according to the manufacturer’s protocol, including DNase I treatment (EURx). RNA concentration and purity were assessed using a NanoPhotometer® N60 (IMPLEN). Samples with A260/A280 and A260/A230 absorbance ratios greater than 2.2 were used for further analysis. RNA integrity was assessed by electrophoresis on a 1% native agarose gel.

Single-stranded cDNA was synthesized from 2 µg of total RNA in a final reaction volume of 10 µL using the Maxima First Strand cDNA Synthesis Kit for RT–qPCR (Thermo Scientific), according to the manufacturer’s instructions. The cDNA was stored at −20 °C until use.

RNA extraction and cDNA synthesis were performed using three biological replicates for each sample group.

#### Real-time quantitative PCR analysis

Quantitative PCR (qPCR) was performed using Power SYBR Green Master Mix (Applied Biosystems). The reaction program consisted of (1) UDG activation at 50 °C for 2 min, (2) Dual-Lock DNA polymerase activation at 95 °C for 2 min, and (3) 40 cycles of denaturation at 95 °C for 1 s and annealing/extension at 60 °C for 30 s. Product specificity was verified by melting-curve analysis and agarose gel electrophoresis. Reactions were run on a QuantStudio™ 5 Real-Time PCR System. Primer sequences are listed in Supplementary Table 1.

For each well, the cycle threshold (Ct) was calculated using QuantStudio Design and Analysis software. Relative transcript levels were calculated using the comparative Ct method and normalized to the most stable reference gene in the dataset. Differences between the N and HS groups were evaluated using Student’s t-test.

### Photocrosslinking and ³²P end-labeling of co-immunoprecipitated nucleic acids

#### BrdU incorporation into Kc cells

For 5-bromo-2-deoxyuridine (BrdU) incorporation, 200 mL of exponentially growing Kc cells (1.2–1.8 × 10 cells/mL) were incubated with 20 µM BrdU for a further 26–27 h. Incorporation was quantified using an anti-BrdU monoclonal antibody, as described previously [17]. 5-Bromo-2-deoxycytidine (BrdC) incorporation was performed similarly.

Cell fractionation of ³²P-labeled and UV-crosslinked cells

Cell fractionation was performed as described by Rzepecki and Fisher [22]; see also Meller et al. [20]. Briefly, cells were collected by centrifugation and washed with PBS. Lysates were prepared with a Dounce homogenizer, filtered, and centrifuged at 10,000 × g for 10 min to obtain the P-10 pellet and S-10 supernatant. The resulting supernatant was then centrifuged at 13,000 × g for 60 min to obtain the P-130 fraction.

#### In vivo photocrosslinking

In vivo photocrosslinking was performed as described previously [17, 22], using a handheld UV lamp (UVGL-58, UVP Inc., San Gabriel, CA) emitting light at 366 nm. All quantities refer to an initial culture volume of 200 mL. For experiments with distamycin and chromomycin, cultures were supplemented with 50 µM distamycin A3 and/or chromomycin (Sigma) 24 h before collection. For heat shock experiments, cells were assigned to three groups (N, HS, and recovery [R]), treated as described above, and then harvested, washed, and resuspended in PBS for photocrosslinking.

Harvested cells were washed, resuspended in 8 mL of PBS, and exposed to 366-nm UV light as described previously.

#### Immunoprecipitation and ³²P end-labeling of co-immunoprecipitated nucleic acids

The procedure was described previously [17, 22]. Following in vivo photocrosslinking, cells were either fractionated or lysed directly, and the resulting material was denatured by adding SDS and DTT to final concentrations of 5% and 20 mM, respectively, followed by boiling for 10 min. Typically, approximately 6 × 10 cells were collected, resuspended in 1.5 mL of PBS, and lysed. The lysates were either denatured immediately or fractionated and then denatured by adding a solution containing 10% SDS and 40 mM DTT. Whole-cell lysates and subcellular fractions were stored at −20 °C until use. Samples derived from approximately 1.5 × 10 cells each were thawed by boiling for 10 min and cooled to RT. Trichloroacetic acid (TCA) was added to a final concentration of 10%, and samples were incubated for 10 min at RT. The precipitated proteins were pelleted by centrifugation at 10,000 × g for 10 min at RT. After the supernatant was removed, 5 µL of 1.5 M Tris-HCl (pH 8.8) was added, and the protein pellet was dissolved in 200 µL of Buffer IPA (20 mM Tris-HCl [pH 8.0], 150 mM NaCl, 0.1% Triton X-100, and 0.02% SDS). Immediately before use, Buffer IPA was supplemented with 2.5 mM CaCl, 0.5 mM PMSF, 1 mM TPCK, 1 mM TLCK, 1 µg/mL leupeptin, and 1 µg/mL pepstatin. Unless otherwise indicated, 5 µL of micrococcal nuclease (680 µg/mL; Boehringer) was added, and samples were incubated at 37 °C for 60 min.

Immunoprecipitation was performed using a modification of the procedure described by Rzepecki and Fisher [22]. Protein A–Sepharose beads (Pharmacia, Piscataway, NJ) were incubated with purified IgG (2–4 µg per 50 µL of beads) at 37 °C for 90 min. After the unbound antibody solution was removed, the beads were incubated with the prepared lysates at 37 °C for 90 min. Bead-bound immunocomplexes were collected by centrifugation and washed three times with 350 µL of Buffer IPB, followed by three washes with 350 µL of Buffer IPA [17, 10].

After the final wash, immunoprecipitates were resuspended in 80 µL of 1.5× T4 polynucleotide kinase buffer (New England Biolabs, Beverly, MA). Next, 1–2 µL of [γ-³²P]ATP (4,500 Ci/mmol; 10 µCi/µL; ICN Pharmaceuticals, Costa Mesa, CA) and 5 units of T4 polynucleotide kinase (New England Biolabs, Beverly, MA) were added. No unlabeled ATP was added. The immunoprecipitates were labeled with ³²P for 30 min at 37 °C, washed three times with 350 µL of Buffer IPB, and resuspended in 45 µL of 2.5× standard SDS–PAGE loading buffer. Samples were separated on a 7% SDS–polyacrylamide gel and transferred to nitrocellulose for autoradiography, phosphorimager analysis (Molecular Dynamics 445 SI), and immunoblotting.

#### Nuclease treatment

Nuclease digestion was performed as described previously [17]. To assess the nuclease sensitivity of ³²P-labeled nucleic acids photocrosslinked to Top2, samples were incubated for 30 min at 37 °C with DNase I (66 µg/mL) or RNase A (60 µg/mL). Digestions were performed in Buffer IPA (20 mM Tris-HCl [pH 8.0], 150 mM NaCl, 0.1% [v/v] Triton X-100, and 0.02% [w/v] SDS) supplemented with 5 mM MgCl and 2.5 mM CaCl.

#### Quantification by scanning densitometry

Unless otherwise indicated, immunoblots and autoradiograms (or phosphorimager scans) were quantified using an LKB Ultrascan XL laser densitometer (LKB Instruments Inc., Gaithersburg, MD).

### PFA crosslinking and co-immunoprecipitation

#### Cell collection and crosslinking

Heat shock and cell collection were performed as described for the solubility assay. Cell pellets (25 × 10 cells) were resuspended in 2 mL of 1% paraformaldehyde and incubated for 7 min at RT with gentle rotation. The samples were immediately centrifuged at 1,000 × g for 3 min at RT. Pellets were resuspended in 1 mL of 125 mM glycine to quench the paraformaldehyde, incubated on a rotary evaporator for 5 min, and centrifuged at 4 °C.

Cells were washed with cold PBS supplemented with protease and phosphatase inhibitors, centrifuged, resuspended in lysis buffer (50 mM Tris-Cl [pH 7.5], 150 mM NaCl, 1 mM EDTA, 1% Triton X-100, 0.1% SDS, and Halt Protease and Phosphatase Inhibitor Cocktail [Thermo Scientific]), and transferred to 1.5-mL tubes. Samples were mechanically homogenized with plastic pestles and incubated on ice for 20 min. The lysates were sonicated using a Bandelin Sonoplus Mini 20 for 25 min at 90% amplitude in 30-s on/off cycles. Sonication conditions were optimized to produce DNA fragments shorter than 500 bp without detectable protein loss. The lysates were then centrifuged at 8,000 × g for 10 min at 4 °C, and the supernatants were collected for further analysis.

#### Co-immunoprecipitation

For each sample, 25 µL of Pierce Protein A/G Magnetic Beads (Thermo Scientific #88802) were used according to the manufacturer’s instructions. Initially, 200 µL of cell lysate was combined with 10 µg of primary antibody (anti-lamin Dm or anti-Top2), and the volume was adjusted to 500 µL with lysis buffer. Samples were incubated with rotation for 1.5 h at RT, transferred to prewashed beads, and incubated for a further 1 h at RT with rotation. Beads were collected using a magnetic stand (Thermo Fisher #CS15000) between washing steps. The flow-through fraction, containing excess unbound antibodies and lysate, was collected before the first wash. Bound material was eluted by adding 100 µL of Laemmli sample buffer and heating at 96 °C for 10 min. After cooling, the supernatants were collected.

Negative controls were processed identically, except that primary antibodies were omitted. Normal polyclonal rabbit IgG (R&D Systems #AB-105-C) was used as an isotype control.

For each sample, 10 µL was loaded onto a polyacrylamide gel alongside the corresponding controls (input, negative control, and flow-through fractions). Proteins were detected by immunoblotting using primary antibodies against lamin Dm, Top2, and lamin C (LC28.26), followed by HRP-conjugated anti-rabbit or anti-mouse IgG, as described in the Antibodies section.

#### Western blot quantification of protein levels

Western blot membranes were imaged using Image Lab software, version 5.2.1 (Bio-Rad Laboratories). Band densitometry was performed using the Volume Analysis tool. Identically sized rectangular regions of interest were defined for each band. Quantification was based on the Mean Value (Int), representing the mean pixel intensity within the selected region.

For protein quantification, each membrane contained a molecular-weight marker and samples from three independent biological replicates. Each replicate included four conditions: N (control), HS (heat shock), R (recovery), and NR (recovery control). To detect proteins of different molecular weights on the same membrane, membranes were cut horizontally into strips corresponding to the expected molecular-weight ranges of the target proteins. Each strip was then incubated with the appropriate antibodies, allowing different proteins to be analyzed from the same set of samples.

At least three technical replicates per protein were analyzed on separate membranes (n = 9). For analyses involving 12 measurements (N = 12), data were obtained from three independent biological experiments, each measured in four replicates. If staining was inadequate for a band or membrane region corresponding to a particular biological replicate, the affected measurement was excluded from the relevant technical replicate.

For each protein, densitometric values were normalized in two steps:

Normalization to the housekeeping protein. The Mean Value (Int) of the target protein was divided by the corresponding Mean Value (Int) of α-tubulin in the same lane to account for differences in sample loading and transfer efficiency.

Normalization to the mean of the biological replicate. Each α-tubulin-normalized value was divided by the mean of the four experimental conditions (N, HS, R, and NR) within the same biological replicate. This step accounted for differences between membranes and allowed relative protein levels to be compared across conditions.

Normalized data were analyzed in GraphPad Prism 10.6.1. Differences between experimental conditions were assessed using ordinary one-way analysis of variance (ANOVA). Statistical significance was defined as p < 0.05; nonsignificant results were reported as ns (p > 0.05).

For the solubility assay, each fraction (lysate, 50 mM, 150 mM, and 500 mM NaCl) was assessed in three biological replicates, with at least three technical repetitions. The three biological samples for each fraction were loaded on the same gel. Band intensities were normalized to the mean densitometric signal within the corresponding fraction. Normalized values were averaged separately for N and HS samples, and standard deviations were calculated. Because fractions were quantified separately, comparisons were restricted to N versus HS within each fraction. Here, a biological replicate denotes a separately cultured cell population subjected to an independent fractionation, whereas a technical replicate denotes a repeated Western blot measurement of the same biological sample.

#### Statistical analysis and data presentation

Statistical analyses (Fisher’s exact test and Student’s t-test, unless otherwise specified) were performed in Microsoft Excel. Quantitative data were plotted using GraphPad Prism 8.0. The number of replicates and the statistical tests used are described in the corresponding figure legends. Independent experiments reported in figure legends were treated as biological replicates.

## 3. Results

### Induction of heat shock and recovery in *Drosophila melanogaster* cultured cells and embryos

We investigated the heat shock (HS) response in Drosophila Kc and S2 cells and, where indicated, in embryos and larvae at the appropriate stages. Both cell lines express lamin Dm, whereas lamin C protein is detected in Kc cells. Hsp70 immunostaining was used to monitor HS induction and recovery (R). After HS, Hsp70 staining increased in Kc and S2 cells (Figure 1A,B). In a subset of heat-shocked cells, Top2 staining became granular or showed an altered sublaminar pattern; some nuclei contained one or several prominent Top2-positive granules. These changes were observed in both cell lines and became less apparent during recovery (Figure 1A,B).

**Figure 1.**
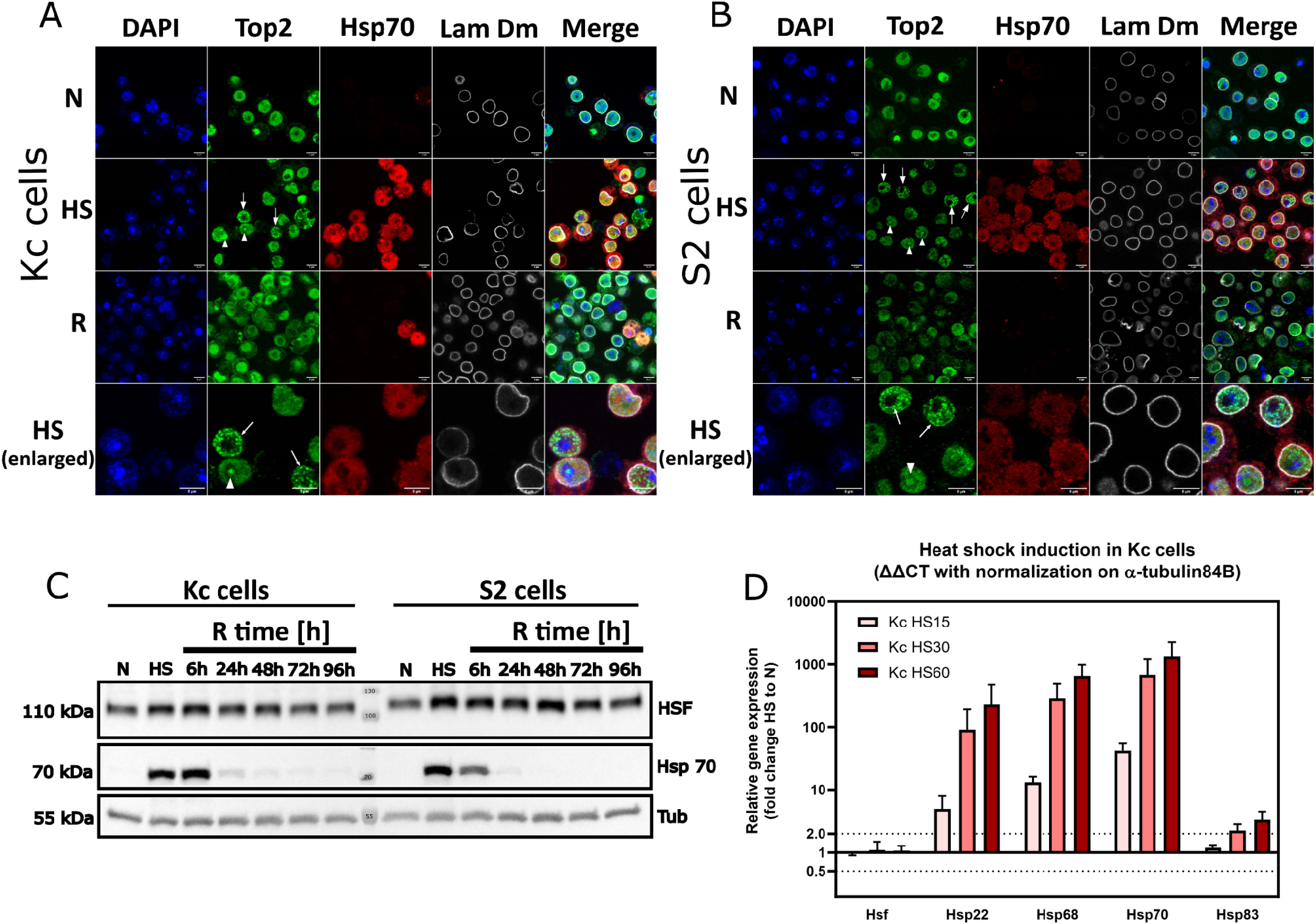
Analysis of heat-inducible Hsp70 expression provides an indicator of the efficiency of the stress response and recovery. (A) Immunofluorescence staining of Hsp70 protein in Kc cells under normal (N), heat shock (HS, 1 h at 37 °C), and recovery (R) conditions. Recovery duration was 72 h. Fluorescence channels: blue – DAPI (nuclei), green – Top2, red – Hsp70, gray – lamin Dm, and merge. Arrows point to cell nuclei with typical Top2 multigranular phenotype. Arrowheads point to cell nuclei with Top2 phenotype of one or a few larger granules. (B) Immunofluorescence staining of Hsp70 in S2 cells under normal (N), heat shock (HS, 1 h at 37 °C), and recovery (R) conditions. Recovery duration was 48 h. Fluorescence channels: blue – DAPI (nuclei), green – Top2, red – Hsp70, gray – lamin Dm, and merge. Arrows and arrowheads as in A. (C) Western blot analysis of Hsp70 protein levels in Kc and S2 cells under heat shock and recovery conditions. The efficiency of recovery was assessed by monitoring the disappearance of the Hsp70 band over time. In addition to Hsp70, HSF was used as a stress response marker, and α-tubulin (α-Tub) served as a loading control. Protein levels were verified at multiple time points following 1 h of heat shock and during recovery at 6, 24, 48, 72, and 96 h. Normal (N) conditions were used as a reference for complete recovery. (D) RT-qPCR analysis of the relative expression levels of selected transcripts involved in the heat shock response, including heat shock proteins (Hsp22, Hsp68, Hsp70, and Hsp83) and the transcriptional regulator Hsf, in Kc cells at different heat shock time points (15, 30, and 60 min). Fold changes were calculated using the ΔΔCt method, with normalization to control cells (N) and the endogenous reference gene Tub84B. Data represent the mean ± standard deviation from two biologically independent experiments, each performed with three technical replicates, except for two cases where only two technical replicates were available due to experimental limitations. qPCR reactions were performed using the QuantStudio™ 5 thermocycler, and data were analyzed using the Applied Biosystems™ qPCR analysis module, with additional calculations performed in Microsoft Excel. Final graphs were generated using GraphPad Prism 10.6.1.

Western blot analysis showed that the Hsp70 signal decreased more rapidly during recovery in S2 cells than in Kc cells (Figure 1C). Most of the Hsp70 signal in S2 cells disappeared within 6 h, whereas Kc cells retained substantial Hsp70 levels at that time. In both cell lines, the Hsp70 signal declined by more than 95% within 24 h. Embryos also exhibited reduced Hsp70 immunofluorescence after 4 h of recovery (Supplementary Figure 1). RT-qPCR analysis detected induction of HS-responsive transcripts, including Hsp70, in Kc cells after 15, 30 and 60 min of HS (Figure 1D). Thus, the Hsp70 response was transient under the conditions examined.

However, these measurements alone do not provide data on complete recovery of every nuclear protein or cellular function.

Based on the Hsp70 time course, we selected 1 h of HS and recovery periods of 48 h for S2 cells and 72 h for Kc cells for subsequent experiments, unless otherwise indicated (Figure 1C). Because HS may affect housekeeping-gene expression, candidate RT-qPCR reference transcripts were evaluated using geNorm. Ribosomal RNAs, actin5C and tubulin84B showed relatively stable expression in the comparative analyses (Supplementary Figure 2B).

Based on these data, 18S/28S ribosomal RNA, actin5C and tubulin84B were selected as RT-qPCR references. Lamin C transcripts were detected in S2 cells at least seven amplification cycles later than in Kc cells (Supplementary Figure 2A). These transcript data do not, by themselves, establish the presence or absence of lamin C protein in S2 cells but are consistent with published reports of nearly undetectable lamin C protein in S2 cells [31, 32]. Please note that data shown are calculated without normalization.

The RT-qPCR comparisons of lamin Dm, lamin C, Top2 and Hsf did not identify statistically significant HS-dependent changes in transcript abundance under the conditions analyzed (Figure 2A). Note the increasing trend for Hsf in both cell lines and opposing trends for LamC; these differences were not statistically significant. Responses of selected HS-inducible transcripts are shown in Figure 2B.

**Figure 2.**
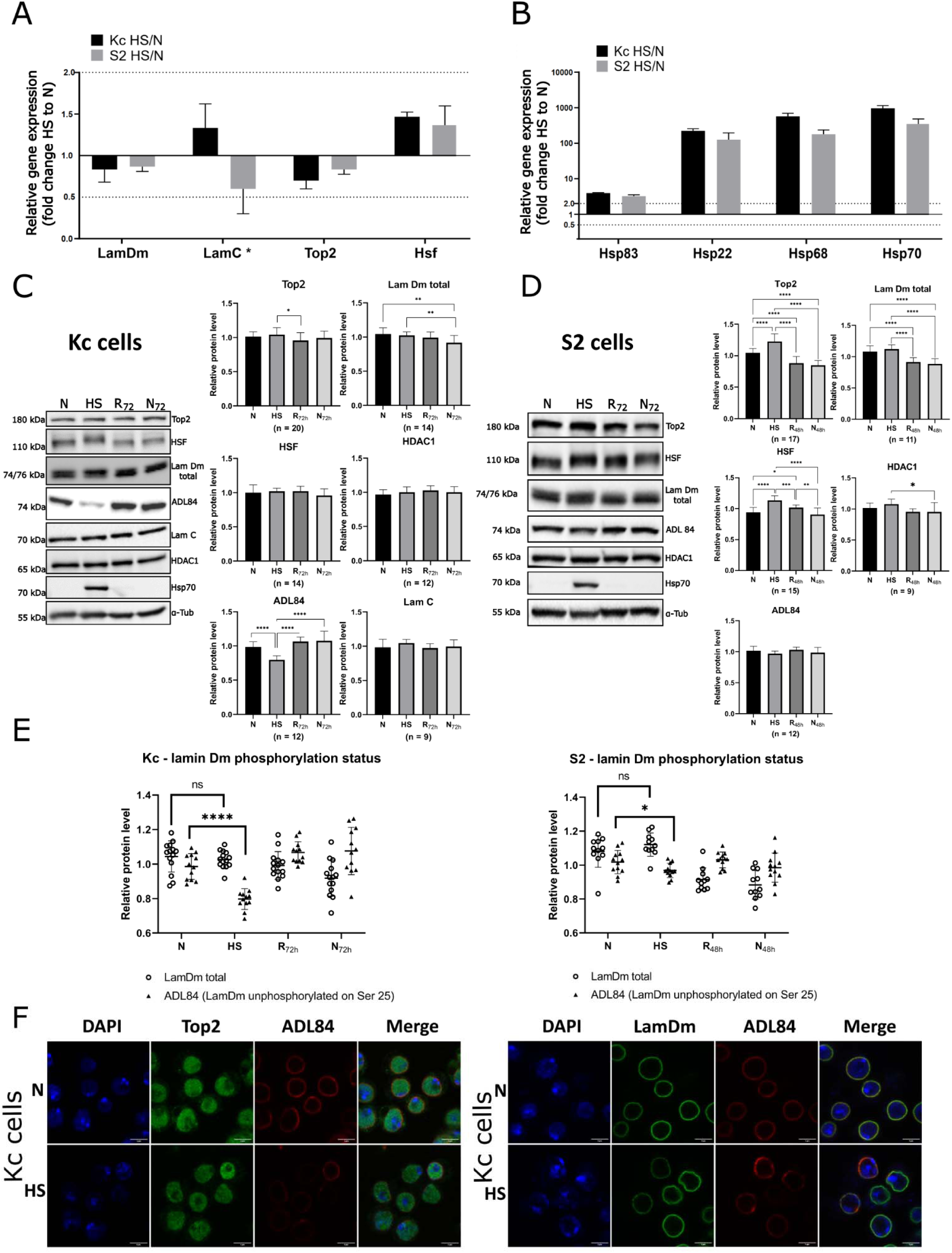
Effects of heat stress on gene and protein expression, and phosphorylation status of lamin Dm, in *Drosophila* cell lines. **(A)** Relative mRNA expression levels of *LamDm*, *LamC*, *Top2*, and *Hsf* genes in Kc (black bars) and S2 (gray bars) cells following 1 hour of heat shock (HS) induction at 37°C. **(B)** Relative mRNA expression of *Hsp83*, *Hsp22*, *Hsp68*, and *Hsp70* genes under the same conditions. Transcript levels were quantified using the Applied Biosystems™ qPCR analysis module in conjunction with a QuantStudio™ 5 thermocycler. Fold changes were calculated using the ΔΔCt method, normalized to non-heat-shocked control cells (N) and a panel of endogenous reference genes (*18S*, *28S*, *Tub84B*, *Act5C*). Error bars represent the standard deviation from three independent biological replicates. Statistical significance was assessed using the unpaired two-tailed t-Student test. A dotted line indicates a 2-fold change threshold on the graph. *#Note:* In Panel A, *LamC* transcripts were detected in S2 cells at approximately the 31st qPCR cycle, about 7 cycles later than in Kc cells, indicating very low expression. This correlates with undetectable level of lamin C protein in S2 cells under these conditions, as assessed by Western blot staining. **(C)** Changes in total protein levels after heat shock induction in Kc cells. Representative Western blot images and corresponding densitometric quantification are shown for Top2, total lamin Dm, Hsf, HDAC1, lamin C, and nonphosphorylated lamin Dm at serine 25 (S25), designated ADL84 (based on antibody specificity). Images for Hsp70 are included as a heat shock (HS) indicator, and α-Tubulin is shown as the loading control. Protein levels were analyzed under four conditions: control (N), heat shock (HS), recovery (R, after 72 h), and time-matched control without HS (NR, 72 h). Densitometry values were normalized first to α-Tubulin and then to the mean intensity value of the biological replicate. Statistical significance between conditions was assessed using ordinary one-way ANOVA (p < 0.05), and significant differences are indicated on the graphs. **(D)** Changes in total protein levels after heat shock induction in S2 cells. Representative Western blot images and corresponding densitometry quantification are shown for Top2, total lamin Dm, Hsf, HDAC1 and nonphosphorylated lamin Dm at serine 25 (S25), designated ADL84. Images for Hsp70 are included as a heat shock (HS) indicator, and α-Tubulin is shown as the loading control. Protein levels were analyzed under four conditions: control (N), heat shock (HS), recovery (R, after 48 h), and time-matched control without HS (NR, 48 h). Densitometric values were normalized first to α-Tubulin and then to the mean intensity value of the biological replicate. Statistical significance between conditions was assessed using ordinary one-way ANOVA (p < 0.05), and significant differences are indicated on the graphs. **(E)** Western blot densitometry analysis of total lamin Dm and unphosphorylated lamin Dm at serine 25 (S25) in Kc and S2 cells. The effect of heat shock on lamin Dm phosphorylation was assessed using ADL84 antibodies specific for the nonphosphorylated form together with rabbit affinity purified anti total lamin Dm antibodies (Lam Dm total). Western blot analysis revealed a reversible decrease in the signal for unphosphorylated lamin Dm after 1 hour of HS, while the total pool of lamin Dm (including both Dm1 and Dm2 forms) remained stable. Densitometry quantification was performed on three biological replicates (technical replicates: total lamin Dm — Kc: n = 14, S2: n = 11; ADL84 — Kc: n = 12, S2: n = 12). Statistical analysis using Student’s t-test showed a significant decrease in unphosphorylated lamin Dm after HS compared to control (Kc: p ≤ 0.0001; S2: p < 0.015), with no significant change in total lamin Dm. Densitometric values were normalized to α-Tubulin and to the mean intensity value of the biological replicate. **(F)** Immunofluorescence analysis of nonphosphorylated lamin Dm at serine 25 (S25) in Kc cells following 1-hour heat shock at 37°C. Using ADL84 antibodies, a decrease in fluorescence signal was observed post-treatment, indicating reduced levels of the nonphosphorylated form of lamin Dm. This supports the Western blot findings and suggests increased phosphorylation of lamin Dm at S25 in response to heat stress. Both images represent Kc cells stained with DAPI (blue) for nuclei. In the left image, Top2 is shown in green and ADL84 in red, and merged overlay. In the right image, total lamin Dm is shown in green and ADL84 in red, and merged overlay. Images were acquired using confocal microscopy.

**Figure 3.**
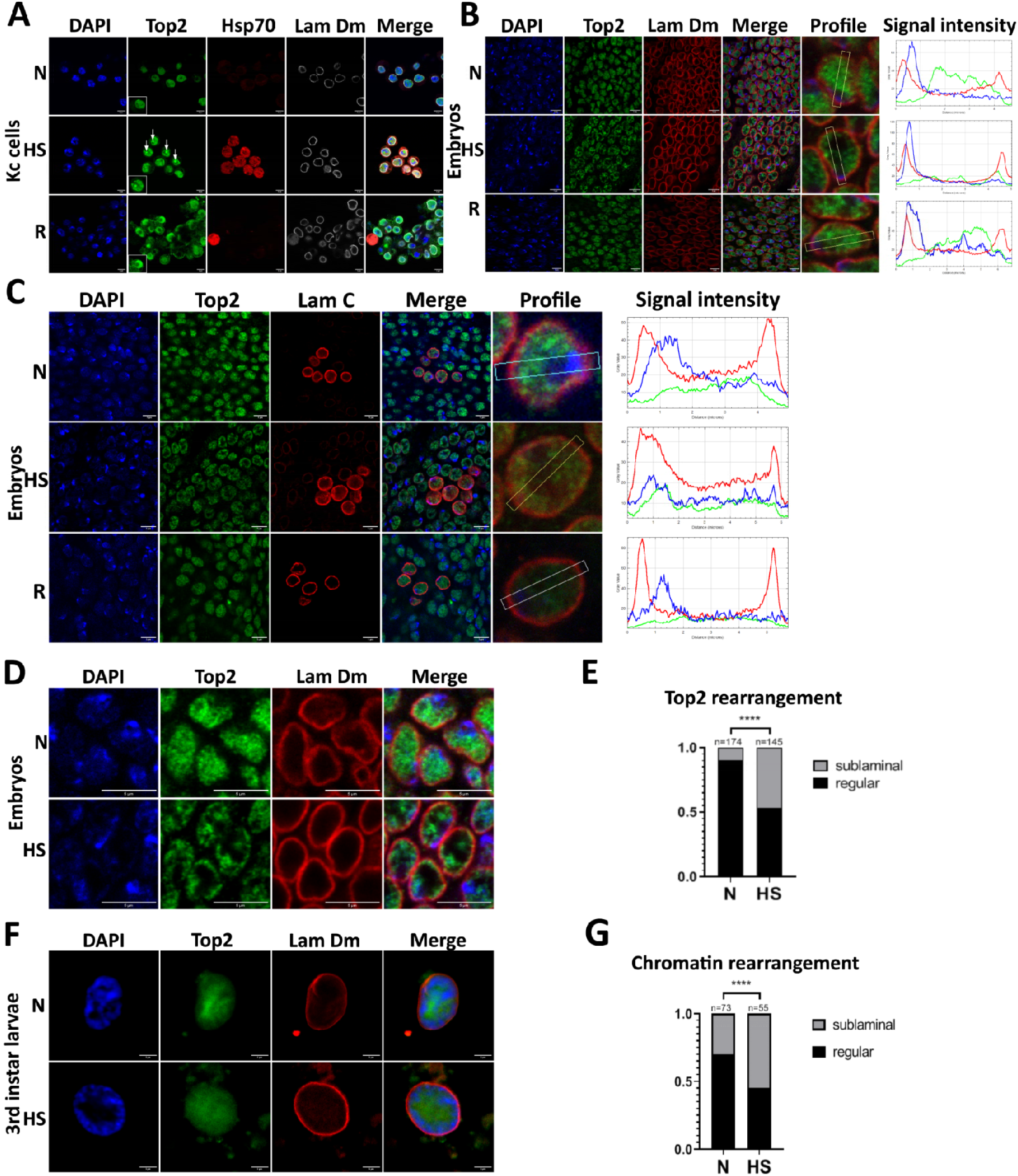
Analysis of the distributions of Top2, lamin Dm, lamin C and DNA in Kc cells and *D. melanogaster* embryos and 3rd larval stage in normal conditions and after induction of heat shock. Immunofluorescence staining of Kc cells (A) and *D. melanogaster* embryos (B) (C), (D), shows stress-induced redistribution and phenotype change of Top2, which is reversible upon recovery. Panel F demonstrates redistribution of chromatin/DAPI (F) upon HS induction in larval polytenic nuclei; Panels E and G demonstrate statistical analyses of D and F respectively. Cells, larvae and embryos were analyzed under N, HS and R conditions. Embryos were from the stage 10-12 as detected from positive signal for lamin C presence in aenocytes. Larvae were at the 3rd stage, nuclei were polyploid with nicely stained chromatin/DAPI. Densitometry graph (when appropriate) with the line density mode view is marked with the line color identical as fluorescent label on IF. **(A)** Immunofluorescence staining of Kc cells for Top2 (green), Hsp70 (red) and lamin Dm (white). DNA stained with DAPI (blue). Arrows point to cell nuclei with typical, single granule/speckle stained for Top2. **(B)** Immunofluorescence staining of embryos (stage10-12) for Top2 (green) and lamin Dm (red). DNA stained with DAPI (blue). **(C)** Immunofluorescence staining of embryos (stage 10-12) for Top2 (green) and lamin C (red). DNA stained with DAPI (blue). **(D)** Immunofluorescence staining of embryos (stage 10-12) for Top2 (green) and lamin Dm (red). DNA stained with DAPI (blue). This panel is a magnified section from Panel B immunofluorescence staining to better illustrate the phenotype. **(E)** Graph demonstrating statistical significance of Top2 relocation to sublaminar region (nuclear lamina vicinity) calculated from immunofluorescence staining for Top2 and lamin Dm in embryos. **(F)** Typical immunofluorescence staining of 3rd larva polyploid nuclei for Top2 (green) and lamin Dm (red). DNA stained with DAPI (blue). Demonstrates two typical phenotypes of distribution of DNA/chromatin: regular and sublaminal (N and HS respectively) for calculation of statistical significance of chromatin relocation. **(G)** Graph demonstrated calculated frequencies of the particular phenotype in polyploid nuclei from immunofluorescence analyses such as in Panel B and D. Top2 rearrangement (E) and chromatin rearrangement (G) after heat shock are presented in the graphs as normalized values (the number of counted cell nuclei marked on the graphs is n). Top2 rearrangement (E) statistics (chi-square test; p=1,9*10E-27) were calculated on the basis of the number of cell nuclei counted in *D. melanogaster* embryonic cells; chromatin rearrangement (G) was calculated on the basis of the number of cell nuclei counted in 3rd instar larval cells (chi-square test; p=6,62*10E-8). Representative staining is shown on the D graph for Top2 rearrangement and on F for chromatin rearrangement.

**Figure 4.**
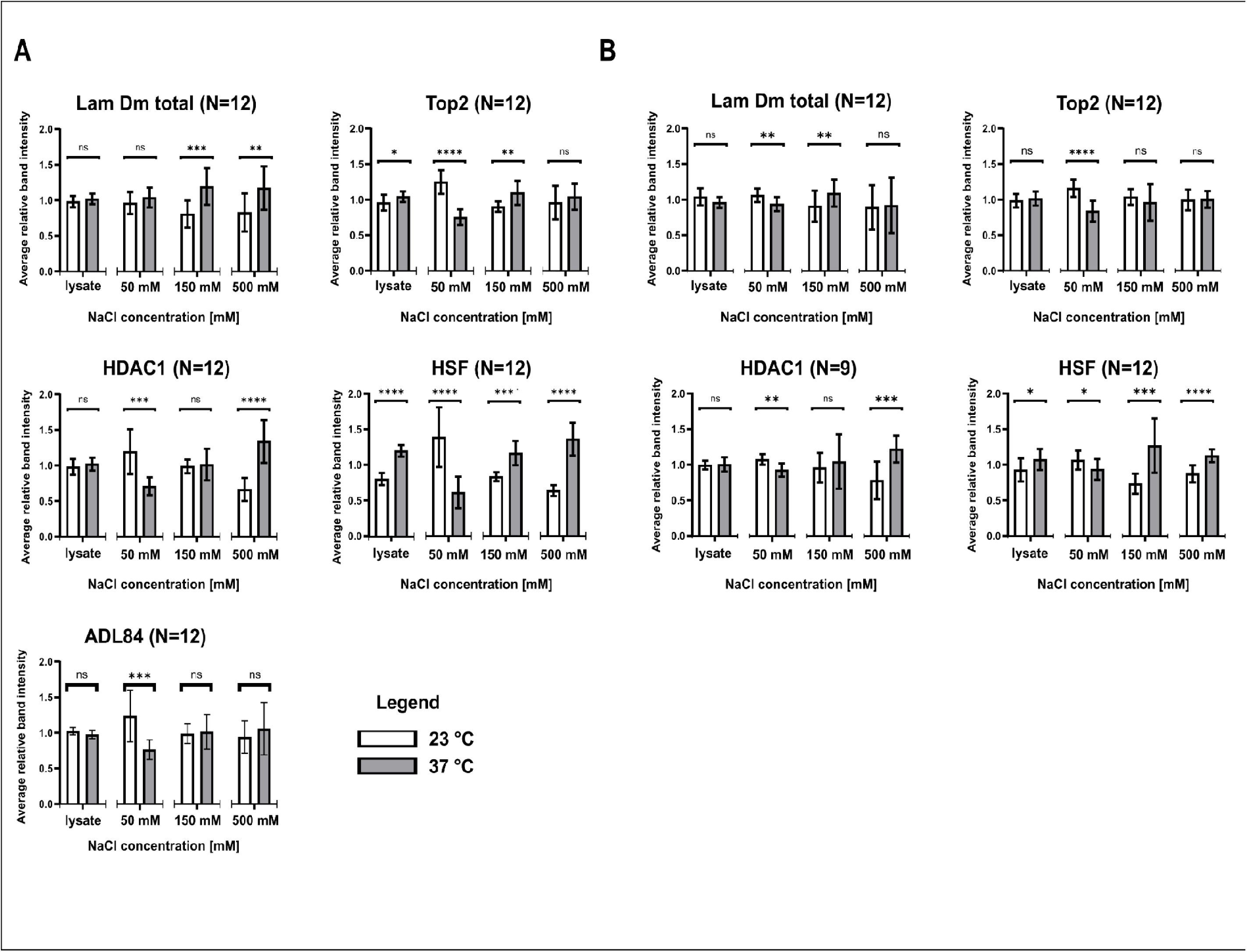
Stress affects the solubility of LamDm and other nuclear proteins in Kc and S2 cells. Statistical analysis of the changes in the solubility of the LamDm, Top2, Hsf and HDAC1 under increasing salt concentrations in Kc cells (Panel A) and S2 cells (Panel B). Control cells and cells subjected to heat shock induction were lysed and extracted sequentially as described in Material and Methods sections using buffer with increasing concentration of NaCl. Low salt buffer contained 50 mM NaCl; Medium salt buffer contained 150 mM NaCl and high salt buffer contained 500 mM NaCl. The amount of the proteins in each fraction was calculated based on Western blots bands intensity using densitometry. Western blots were probed with antibodies for: Top2, HDAC1, HSF and tubulin as a load control. For lamin Dm visualization rabbit affinity purified antibodies were used if total fractions/forms of lamin Dm were to be detected (Lam Dm total) and mouse monoclonal ADL84 antibodies specific for non-phosphorylated S25 for detection of fraction of on phosphorylated S25 lamin Dm fraction. White bars-samples from control cells; Grey bars-samples from heat shock induced cells. Heat shock induction for 60 min at 37oC. The data presented in the bar graphs has been validated for the mean intensity of each band from a given membrane (see M&M section). The statistics (t-student test) on the bar graphs demonstrate the comparison of the N versus HS conditions for each fraction independently. Above each plot, the number of repetitions (usually 3 biological and 4 technical) taken for the statistical analysis is given (N). The white bars in each graph refer to normal conditions (N - 23°C), and the grey bars refer to heat shock induction (HS – 1 h, 37°C). The subsequent fractions are labelled as lysate (control), 50 mM, 150 mM and 500 mM, respectively. The data has been evaluated to identify any significant differences, and these are indicated by the following symbols: “ns” - no significance, * indicates a p value<0,05, ** indicates a p value< 0,01, *** indicates a p value<0,001, and **** indicates a p value<0,0001.

### Heat shock alters lamin Dm electrophoretic forms and S25-sensitive antibody reactivity

We examined the abundance of selected proteins during HS and recovery by Western blotting (Figure 2C,D). In Kc cells, comparisons between normal (N) and HS conditions did not reveal significant changes in the amounts of Top2, lamin Dm, lamin C, HDAC1 or Hsf. In S2 cells, Top2 and Hsf signals increased after HS, whereas the HDAC1 signal did not change significantly in the reported comparison. Hsf also showed altered electrophoretic mobility in both cell lines, consistent with the previously described HS-associated phosphorylation of this protein [33]. In the 48 h recovery and 45 h control-versus-N comparisons, Top2 and lamin Dm signals were significantly lower (Figure 2D). HSF protein levels also differed significantly between all samples except N and N48h. The higher level of HSF compared with the control may result from slower recovery from the elevated HS level.

Western blots obtained with polyclonal anti-lamin Dm antibodies revealed changes in the relative intensities of the two major lamin Dm bands in Kc cells after HS (Figure 2C; see also Figure 5F, G and Supplementary Figure 5). These electrophoretic forms are commonly designated Dm1 and Dm2. The corresponding analysis in S2 cells did not detect a statistically significant HS-associated change in their proportions (Figure 2D). Changes in band mobility or relative intensity are consistent with altered post-translational modification, do not independently identify the responsible phosphorylation site, but are consistent with the previously demonstrated correlation between phosphorylation and migration of lamin Dm bands [13, 3].

**Figure 5.**
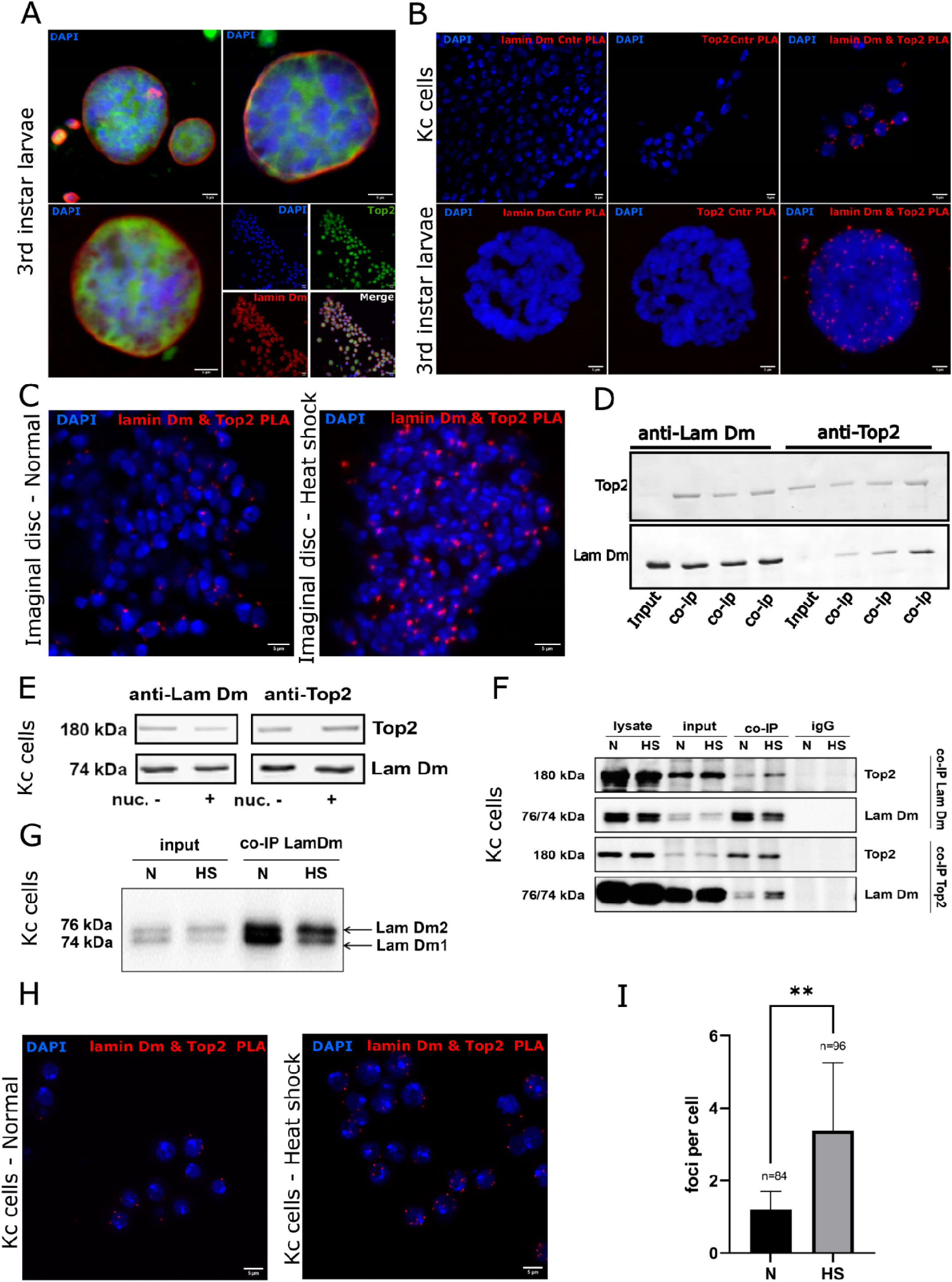
Lamin Dm colocalization and co-immunoprecititation with Top2 protein and their increase during heat shock induction. Panel. **A** Immunofluorescence staining for lamin Dm (ADL67, top left and ADL84 top right) or lamin C (ALC28, bottom left) (red) and Top2 (green) in control conditions in 3rd instar larval tissues. DNA stained with DAPI (blue). For polyploid and polytenic nuclei visualization only merged channels are shown. Bottom, right panels: immunofluorescence staining for diploid cells, split channels with lamin Dm staining (ADL67)(red) and Top2 protein (rabbit polyclonal, affinity purified) (green). **Panel B** Proximity Ligation Assay (PLA) analyses for colocalization of lamin Dm and Top2 in Kc cells (upper panels) and 3rd instar larvae polytenic nuclei of salivary glands (bottom panels). DNA was stained with DAPI. From left to right on both levels: control for anti lamin Dm antibody, control for anti Top2 antibody and PLA signal from both antibodies staining (red) indicates colocalization of proteins. Please see Supplemental Movies for 3D distribution of PLA colocalization signals of lamin Dm and Top2. **Panel C** Proximity Ligation Assay for lamin Dm and Top2 proteins in 3rd instar larvae imaginal discs with the same antibodies as in Panel B. Control larva PLA staining on the left and heat shock induced larva PLA on the right. DNA stained with DAPI. **Panel D and E** Western blot analysis of immunoprecipitation of lamin Dm and anti Top2 from Kc cells extracts prepared under native conditions as described in Materials and Methods section. In Panel E, after IP procedure additional nuclease treatment in beads, and washing steps were added to demonstrate nuclease insensitivity of the co-IP protein. Membrane was divided and stained for lamin Dm with mouse ADL67 antibody (top panel) and for Top2 with rabbit AP purified anti Top2 antibodies (bottom panel). Input-extract sample stained with antibody for lamin Dm or Top2; co-IP-immunoprecipitation anti lamin (or anti Top2) with detected binding (co-IP) partner protein. **(E)** Western blot experiment for additional nuclease in beads treatment and additional washing steps before gel loading. **Panel F and G** Western blot analyses of the immunoprecipitation of lamin Dm and Top2 proteins from control (N) Kc cells and Kc cells with induced heat shock effect (HS). Kc cells were fixed with PFA and extracted under denaturing conditions to ensure efficiency of extractions of major fraction of lamins and Top2 without losing their interaction partners. **Panel G** represents magnification of the part of western blot from **F**. Immunoprecipitation was performed with AP rabbit antibodies for lamin Dm and Top2. Membrane was divided and each strip processed for visualization of signal with the same rabbit antibodies as used for IP. Please note different ratio of both lamin Dm forms in normal Kc cells and heat shocked cells (lysate, input) and the same ratio retained in IP anti lamin Dm (**G**) and as co-IP fraction in anti Top2 immunoprecipitation. Please also note increased bands of co-immunoprecipitated lamin Dm and Top2 from heat shocked Kc cells. **Panel H and I** Proximity Ligation Assay (PLA) analyses for colocalization of lamin Dm and Top2 in control Kc cells and Kc cells with induced heat shock (H), and statistical analyses of induction of additional PLA colocalization signals/loci in heat shocked Kc cells (HS) comparing to control cells (N). Statistical analysis of the Kc PLA (Panel I) revealed a significant increase in PLA signal upon heat shock. Quantification was performed on Kc cells, with approximately 100 nuclei analyzed per condition, and statistical significance was determined using the two-tailed Mann–Whitney U test (*p* < 0.01).

To examine whether HS was associated with changes at the lamin Dm S25 epitope, we used the monoclonal antibody ADL84, which recognizes the nonphosphorylated S25-containing region of lamin Dm [34, 15]. In Kc-cell immunoblots, ADL84 reactivity strongly decreased after HS and increased during recovery which correlated with changes in the intensities of the lower and upper lamin Dm bands in the polyclonal immunoblot (Figure 2C). The corresponding comparison in Figure 2D showed no changes in S2 cells, whereas the separate analysis presented in Figure 2E indicated reduced ADL84 reactivity after HS (see also immunoblots with polyclonal antibodies and ADL84 in Supplemental Figure 5). Because ADL84 is phosphorylation-sensitive, reduced binding is consistent with increased S25 phosphorylation. The calculated, statistically significant average increase in S25 phosphorylation in both experiments may seem a little low but indicates that about 20% of lamin Dm1 molecules were phosphorylated to form lamin Dm2 overall (phosphorylation/dephosphorylation equilibrium upon stress induction. Taking into account preexisting lamin Dm molecules already phosphorylated on S25 and population of lamin Dm highly polymerized in filaments and interacting with proteins with no accessible S25 phosphorylation site makes the numbers physiologically important. See the Discussion for further considerations.

Immunofluorescence staining with ADL84 was reduced in Kc cells after HS (Figure 2F). Changes in fluorescence intensity should not be treated as a direct measurement of S25 phosphorylation level but only as an additional indicator. Note the greater irregularity of ADL84 staining compared with polyclonal immunostaining for lamin Dm. The magnitude of the ADL84 response and the abundance of selected proteins differed between Kc and S2 cells (Figure 2C–F). In contrast, both cell lines showed HS-associated alterations in Top2 nuclear staining, including the appearance of Top2-positive granules (Figure 1A,B).

### Heat shock alters Top2 localization in cultured cells and embryos and chromatin distribution in larvae

We next examined the subnuclear distribution of Top2 and lamins by immunofluorescence. In Kc and S2 cells, HS was associated with a shift from predominantly diffuse nuclear Top2 staining to patterns that included intranuclear granules and altered peripheral or sublaminar staining. Figure 3A also illustrates additional phenotypes compared with those in Figure 1A,B. In stage 10–12 embryos, HS increased the proportion of nuclei displaying Top2 enrichment near the nuclear periphery (Figure 3B,C). The embryonic HS-induced phenotype of lamina-associated Top2 is significantly different from the prominent Top2-positive granules observed in the cultured cell lines. No obvious redistribution of lamin Dm or lamin C was detected in the analyzed preparations (Figure 3A–C). The altered Top2 staining patterns became less apparent after recovery.

Quantification of embryonic immunofluorescence images confirmed a significant increase in nuclei with peripheral Top2 staining after HS (Figure 3D,E). In a separate experiment, DAPI staining of third-instar larval nuclei revealed a significant increase in the proportion of nuclei with chromatin concentrated near the nuclear periphery (Figure 3F,G). The shift of chromatin toward the nuclear lamina was not associated with concomitant shift of Top2 relocation to nuclear lamina as is the case of embryonic cells and Kc cells. Nevertheless, Top2 was more uniformly distributed after HS (Figure 3F,G).

Additional immunofluorescence analyses are presented in Supplementary Figures 3 and 4. In Kc cells, Hsp90 displayed expected increased nuclear staining after HS (Supplementary Figure 3A), whereas no obvious redistribution of lamin Dm, HDAC1, LBR, bocksbeutel or RNA polymerase II phosphorylated at Ser5 was observed in the preparations examined (Supplementary Figure 3B–D). Embryonic staining for HP1, HSF, HDAC1 and phosphorylated RNA polymerase II is shown in Supplementary Figure 4. The data do not demonstrate any changes in distribution or intensity.

### Heat shock changes the salt extractability of lamin Dm, Top2, HDAC1 and HSF

We compared the sequential salt extractability of lamin Dm, Top2, HDAC1 and HSF in control and heat-shocked Kc and S2 cells (Figure 4A,B; representative sample blots and triplicates are shown in Supplementary Figure 5). Proteins were extracted using buffers containing 50, 150 and 500 mM NaCl, as described in Materials and Methods. Band intensities were quantified separately for each fraction, and N-versus-HS comparisons were made within the same extraction condition. Changes in extractability do not necessarily indicate changes in total cellular protein abundance or prove redistribution within a specific nuclear location but should reflect HS-induced changes in protein interactions and/or properties.

In Kc cells, HS increased the amount of lamin Dm recovered in the 150 and 500 mM NaCl fractions (Figure 4A). Top2 was reduced in the 50 mM fraction and increased in the 150 mM fraction. HDAC1 was reduced in the 50 mM fraction and increased in the 500 mM fraction, whereas HSF was reduced in the 50 mM fraction and increased in the higher-salt fractions; an increase in the whole-cell HSF signal was also observed. The ADL84-reactive lamin Dm fraction was reduced in the 50 mM NaCl extract after HS (see also Supplementary Figure 5A). This result identifies a change in the extraction behavior of an antibody-defined lamin Dm population but does not independently establish a particular Dm1-to-Dm2 conversion but, when analyzed together with the data in Supplementary Figure 5, it is evident that the conversion of Dm1-to-Dm2 takes place (Lam Dm total) and less lamin Dm1 is detected with ADL84 antibody.

S2 cells showed a partly different extraction profile (Figure 4B). Total lamin Dm soluble fraction was reduced in the 50 mM NaCl fraction and increased in the 150 mM fraction. HDAC1 was reduced in the 50 mM fraction and increased in the 500 mM fraction (similar to Kc cells), while HSF was reduced in the 50 mM fraction and increased in the higher-salt fractions - the same way as in Kc cells. Changes in Top2 extractability were also observed in S2 cells (Figure 4B). Overall, these experiments indicate HS-associated, protein-and cell-line-dependent changes in salt extractability rather than a uniform redistribution of all tested proteins. Nevertheless, HS-related changes for HDAC1 and HSF were the same in both cell lines, and no significant contradictory changes were detected. Some protein fractions in Kc cells were significantly modified when no changes were detected in S2 cells and vice versa. For example, the Top2 fractions extracted with 50 and 150 mM NaCl changed in Kc cells whereas only the 50 mM fraction changed in S2 cells, with no changes detected in the 150 mM fraction.

Representative immunoblots illustrating the fractionation patterns and changes in lamin Dm electrophoretic forms are provided in Supplementary Figure 5A,B.

### Lamin Dm and Top2 show proximity and co-immunoprecipitation; heat shock increases a nucleic acid-associated signal

Earlier studies identified *in vivo* associations of lamin Dm with nucleic acids [17] and characterized nucleic acid-associated Top2 populations [22]. Here, immunofluorescence indicated that lamin Dm was predominantly peripheral, whereas Top2 was distributed mainly within the nuclear interior (Figure 5A). Proximity ligation assays (PLA) detected lamin Dm–Top2 proximity signals in Kc cells and in polytene nuclei of third-instar larval salivary glands (Figure 5B). Single-primary-antibody controls did not produce comparable signals. Numerous foci of different sizes were detected in polytene nuclei of salivary glands, whereas substantially fewer foci were seen in diploid cells. Three-dimensional imaging placed many signalsl near the boundary between polytene chromosomes and the nuclear lamina, with others in the nuclear interior (Supplementary Movies 1 and 2). PLA signals also appeared more numerous or prominent after HS in imaginal-disc preparations (Figure 5C) and Kc cells (Figure 5H); the reported increase in mean foci per nucleus was significant in Kc cells (Figure 5I). PLA detects molecular proximity within the assay’s spatial range; it does not by itself demonstrate direct protein–protein binding or an association with lamina-associated domains. The number of proximity foci in salivary gland nuclei and diploid cells may reflect the small population of lamin Dm (roughly 20%) being phosphorylated upon HS, lack of relocation of chromatin in Kc, S2 and in embryonic cells compared with polyploid/polytenic nuclei, high polyploidy of salivary gland nuclei with multiplied potential foci or specific structure of chromatin in polytenic nuclei. Additional explanations may apply to imaginal disc cells as a quiescent stem-cell population.

To further examine the association between lamin Dm and Top2, we performed co-immunoprecipitation (co-IP) from native Kc-cell extracts using affinity-purified antibodies against either protein (Figure 5D). Top2 and lamin Dm were detected in the corresponding co-immunoprecipitates. Additional on-bead S7 nuclease treatment did not visibly abolish co-IP under these native conditions (Figure 5E). These results support the presence of lamin Dm–Top2-containing complexes extracted under native conditions but do not establish whether their association is direct or mediated by other proteins or residual nucleic acids or both.

Because only a small proportion of lamin Dm could be extracted under the native conditions, we also analyzed Kc cells fixed with paraformaldehyde before extraction and sonication. The lysis buffer contained 150 mM NaCl, 1% Triton X-100 and 0.1% SDS, and chromatin was sheared to fragments of approximately 500 bp, as described in Materials and Methods. Control and heat-shocked cells were processed in parallel to compare the recovered complexes.

Reciprocal immunoprecipitation after paraformaldehyde crosslinking recovered lamin Dm and Top2 in both N and HS samples (Figure 5F). The lamin Dm band pattern in the Top2 co-immunoprecipitate resembled that in the corresponding input and lamin Dm immunoprecipitate (Figure 5F,G). In both IPs the co-isolated protein band appears stronger in HS then N which suggest higher amounts of complexes or higher extractability of protein complexes under such condition. Therefore, no conclusion may be drawn on protein interactions increase and the potential effect of S25 phosphorylation from this experiment.

We next investigated nucleic acid associations using BrdU incorporation, *in vivo* UV photocrosslinking, immunoprecipitation and, where indicated, ³²P end-labeling of crosslinked nucleic acids, followed by immunoblotting and autoradiography (Figure 6; [17, 22]). The procedure is described in Materials and Methods.

**Figure 6.**
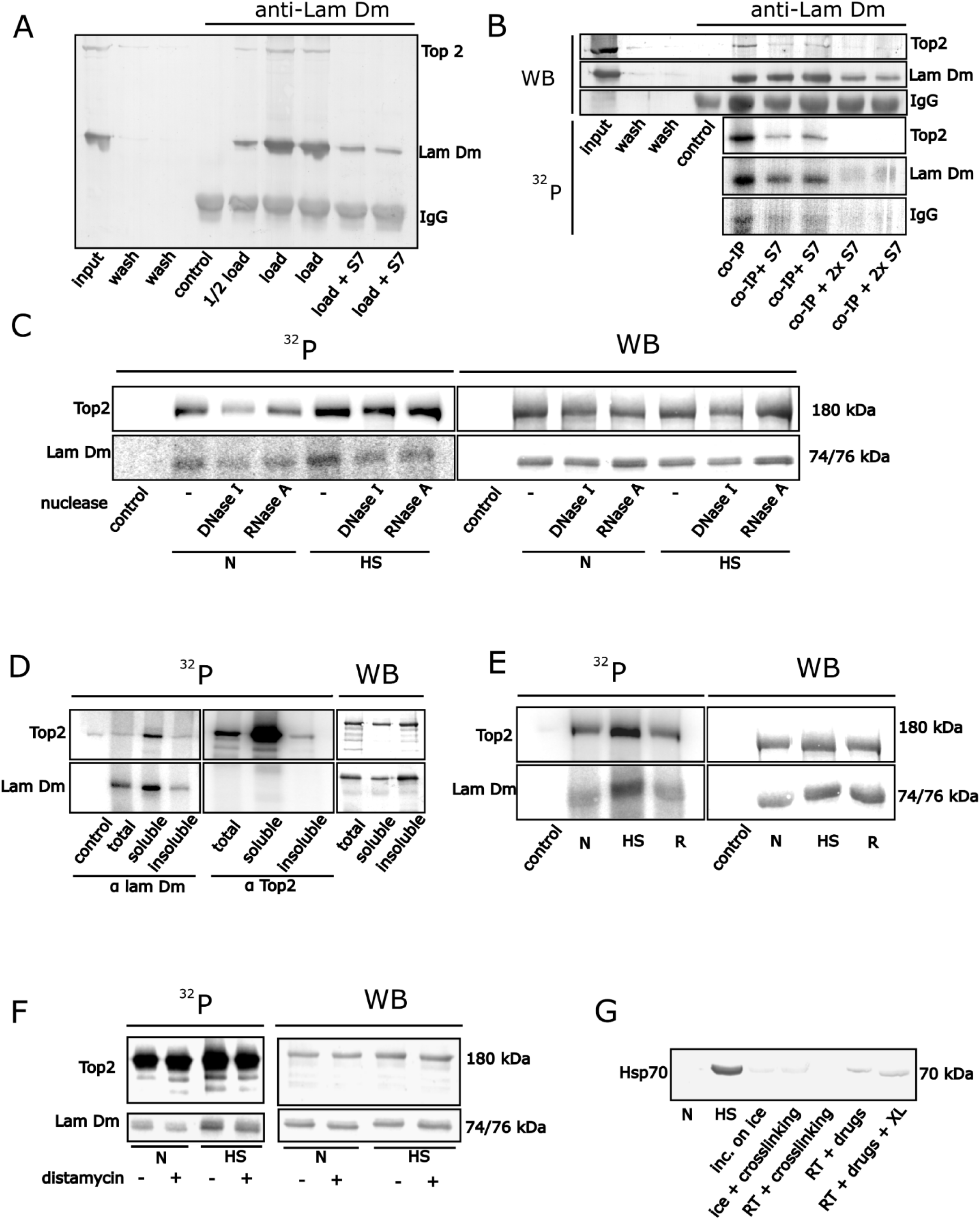
Control Kc cells were used alone for the experiments in A, B and D and together with heat shock induced Kc cells in C, E and F. Immunoprecipitation was with AP rabbit antibodies for lamin Dm and for Top2. Proteins visualization was typically with ADL67 for lamin Dm and with rabbit AP anti Top2 protein with appropriate secondary antibodies and different substrates for lamin Dm and Top2 specific visualization on membranes (see Materials and Methods). WB-western blot analyses; 32P-autoradiography of the nucleic acids radioactive label from labeling with γ32P-ATP and PNK kinase; ½ load/load-half of the normal volume and normal volume of extract loaded onto the beads; +S7-additional S7 nuclease treatment of the beads and additional washing steps added to the procedure; +2xS7-additional, double amount of S7 nuclease with additional washing steps. **Panel A** No radioactive labeling with Poly Nucleotide Kinase (PNK) was used in this experiment. Input-extract to be loaded onto the beads with antibodies; wash-pulled fractions washed off buffer from washing steps; control-IP with isolated IgGs from pre-immuned rabbit serum; ½ load/load-half load and normal volume of extract loaded onto the beads; +S7-additional S7 nuclease treatment of the beads and additional washing steps added to the procedure. Please note the co-IP of Top2 with lamin Dm in this procedure and full nuclease sensitivity of co-immunoprecipitated Top2 band. **Panel B** Western blot analysis of complete crosslinking, IP, radioactive labeling procedure and with similar experimental set up as in A. Please note the co-IP of Top2 band with lamin Dm immunoprecipitation detected in WB and autoradiography and the full sensitivity of this Top2 bands for additional nuclease treatment. Interestingly, also label associated with lamin Dm is also nuclease sensitive up to total disappearance of radioactivity associated with immunoprecipitated lamin Dm band while protein band, despite from the fact of decreasing in intensity still persists. This experiment confirms the co-IP of Top2 with lamin Dm and also confirms that in these BrdU and UV-crosslinking procedure (denaturing conditions) is fully dependent on nucleic acids presence. **Panel C** Western blot analysis of the crosslinking and immunoprecipitation procedure on control Kc cells and cells with induced heat shock with IP anti lamin Dm and IP anti Top2. After entire procedure IP samples were treated with buffer (-/control) or with DNase 1 and RNase A in beads followed by additional washing step and autoradiography and WB analysis. Lanes 2-4: normal cells; Lanes 5-7 heat shocked cells om WB and autoradiography panel. Samples of lamin Dm IP and Top2 IP from heat shocked cells demonstrate higher level of nucleic acids associated with their protein bands but no significant changes in proportions of sensitivity of the label for DNA and RNA digestion was detected in heat shocked cells comparing to control cells. **Panel D** Western blot analysis of the crosslinking and immunoprecipitation procedure on control Kc cells only to test efficiency of nucleic acid binding by total fraction of lamin Dm and Top2 (total) and fractions extracted with medium salt buffer (150 mM NaCl; soluble) and high salt buffer (500 mM NaCl; insoluble). Obtained data suggest that the highest efficiency in nucleic acid binding has lamin Dm and Top2 from “soluble” fraction. **Panel E** Western blot analysis of the crosslinking and immunoprecipitation procedure on control Kc cells and cells with induced heat shock with IP anti lamin Dm and IP anti Top2. Control-control for IP; N-IP and labeling from control Kc cells; HS-IP and labeling from heat shocked cells; R-IP and labeling from heat shocked and “recovered” cells. Data indicate that heat shock transiently increased lamin Dm and Top2 association with nucleic acids. **Panel F** Western blot analysis of the crosslinking and immunoprecipitation procedure on control Kc cells and cells with induced heat shock with IP anti lamin Dm and IP anti Top2. Kc cells were fed with BrdU as in all experiments with addition (+) or not (-) of distamycin and divided into control cells (N) and heat shock cells (HS) before the entire procedure of the UV-crosslinking and labeling. All four-cell population were analyzed in WB and autoradiography. For lamin Dm binding to nucleic acids in control cells we detected lower signal in cells with distamycin as well as in heat shocked cells with distamycin comparing to lack of distamycin. The inhibition of nucleic acid binding of lamin Dm by distamycin agreed with our earlier studies. Lack of changes suggest the same mode of DNA association in N and HS conditions. For Top2 no changes in nucleic acids binding were detected in control cells between no distamycin and with distamycin. In heat shocked cells distamycin decreased the Top2 nucleic acid association might suggest that during heat shock increased binding to DNA was based on minor grove of DNA mostly. **Panel G** Western blot analysis of the control experiment for BrdU and UV-crosslinking procedure. Experiment demonstrates that experimental procedures of BrdU incorporation, incubation on ice, UV crosslinking, incubation with drugs (distamycin, chromomycin) and their combined effect does not induce stress response as judged by hsp70 staining on Western blots.

The assay was applied to Kc cells under control, HS and recovery conditions. Its outcome depends on BrdU incorporation, UV-crosslinking efficiency, protein recovery under denatured conditions and the susceptibility of associated nucleic acids to nuclease treatment. Consequently, it preferentially detects the covalently crosslinked fractions of lamin Dm and Top2 rather than their entire cellular populations.

In the experiment shown in Figure 6A, the radioactive end-labeling step was omitted. Co-immunoprecipitated Top2 was detected with lamin Dm, but its signal decreased after additional S7 nuclease treatment. The Top2 co-IP efficiency was nuclease-dependent and differed from that observed in the native co-IP assay (Figure 5E).

This difference indicates that the detected association depends on the experimental conditions (nucleic acids) and detects only a fraction of the protein populations-those associated with nucleic acids.

In the complete photocrosslinking and ³²P-labeling experiment, both Top2 co-immunoprecipitation and the radiolabeled signals associated with Top2 and lamin Dm decreased with increasing nuclease treatment (Figure 6B). At higher nuclease concentrations, the Top2 band and associated radioactive signal were no longer detectable; lamin Dm recovery and its radioactive signal also decreased. These findings are consistent with the recovery of nucleic acid-associated protein fractions. They do not show directly that the two proteins occupy the same DNA molecule or neighboring sequences.

We then compared nuclease sensitivity of IP for each protein under N and HS conditions (Figure 6C). DNase I and RNase A treatments produced broadly similar patterns of sensitivity, but the nucleic acid-associated Top2 signal was greater in HS samples. The results therefore indicate an increase in the detected nucleic acid-associated fraction of Top2 without establishing a change in the relative contributions of DNA and RNA. Increased PLA proximity signals (Figure 5C,H,I) should be considered separately from the nucleic acid-associated signals obtained with this assay.

When Kc-cell extracts were separated by salt extraction, stronger nucleic acid-associated signals for both lamin Dm and Top2 were detected in the 150 mM NaCl fraction than in the material extracted at 500 mM NaCl (Figure 6D). A radiolabeled Top2-associated band was also detected in material co-immunoprecipitated with lamin Dm, although the corresponding Western blot signal was weak or absent. The comparison of N, HS and recovery samples showed increased nucleic acid-associated signals for limited fractions of lamin Dm and Top2 after HS, followed by a decrease during recovery (Figure 6E).

To explore whether the detected nucleic acid associations were sensitive to an AT-rich-DNA-binding compound, we treated Kc cells with distamycin before the photocrosslinking assay (Figure 6F). Distamycin reduced the radioactive signal associated with lamin Dm in both N and HS samples; its effect on the Top2-associated signal was smaller and depended on the experimental condition. These results demonstrate sensitivity to distamycin treatment but do not identify the bound nucleic acid sequences or prove a change in sequence preference. The experimental procedure alone did not produce detectable Hsp70 induction in the control shown in Figure 6G.

Taken together, the results show that HS-associated changes in ADL84 reactivity and protein extractability were accompanied by increased nucleic acid-associated signals for lamin Dm and Top2 in the photocrosslinking assay. They do not demonstrate that S25 phosphorylation causes the increase in nucleic acid association.

Top2 nuclear staining changed after HS in cultured cells and embryos, whereas peripheral chromatin redistribution was demonstrated separately in third-instar larvae. PLA and co-IP experiments supported the HS associated proximity or association of lamin Dm and Top2, increased number of protein complexes and their coincidence with S25 phosphorylation of lamin Dm fraction. The photocrosslinking experiments detected HS-increased nucleic acid-associated signals in limited protein fraction of Top2 and lamin Dm. These observations should not be interpreted as direct evidence that entire fractions of Top2 and chromatin relocate together or that HS induces tethering of specific genes to the nuclear lamina.

## 4. Discussion

Lamin Dm, chromatin organization and the cellular context of the heat-shock response

The present study addresses how heat shock (HS) changes the properties and associations of lamin Dm and DNA topoisomerase II (Top2), two proteins with established but distinct contributions to nuclear organization. Lamin Dm, like other B-type lamins, participates in organizing peripheral chromatin and interacts with nuclear-envelope and chromatin-associated proteins. Lamina-associated domains (LADs), nucleolus-associated domains (NADs), and topologically associating domains (TADs) provide useful concepts for discussing spatial genome organization, but our experiments did not map any of these domains or their boundaries [35, 36, 37, 38, 39, 2]. Therefore, an alteration in staining, distribution or extractability cannot be taken as evidence that a particular LAD or TAD was formed, dissolved, or moved during HS.

An important experimental distinction between Drosophila lamins is provided by earlier in vivo photocrosslinking studies: interphase lamin Dm was recovered in association with both DNA and RNA, whereas comparable nucleic-acid associations were not detected for lamin C in the same assay [17]. In addition, lamin Dm is not the sole possible link between chromatin and the nuclear envelope: LEM-domain proteins, LBR, BAF, HP1 and nucleoporins provide additional direct or indirect interaction routes [40, 41, 42, 2, 3]. The functional relevance of lamin Dm consequently lies in its participation in a broader interaction network and interaction hub rather than in an exclusive role as a chromatin tether.

### Heat-shock response, recovery and differences between Kc and S2 cells

Hsp70 accumulation was used to verify HS induction and to follow attenuation of the response during recovery. Induction of Hsp70 and HS-responsive transcripts after exposure to 37 °C and relocation of HSP90 to the nuclear compartment confirm activation of a canonical stress response, while the altered electrophoretic mobility of HSF is compatible with the known HS-associated phosphorylation of this transcription factor [33, 43]. A 60-min HS was selected to examine changes after the initial transcriptional response had begun. The Hsp70 signal subsequently declined in both cell lines, more rapidly in S2 than in Kc cells, and was also reduced in embryos after recovery (Figures 1A–D and Supplementary Figure 1). These kinetics indicate that the induced response is transient; they do not, on their own, demonstrate that the transcriptome, proteome, chromatin organization and all nuclear functions have returned fully to their pre-stress state.

Because both cell lines have reported doubling times close to one day, the large decline in the Hsp70 signal within 24 h is unlikely to be explained by dilution during cell division. Protein degradation is the most obvious contributor, although degradation rates were not determined directly. The slower decline in Kc cells may also reflect differences in Hsp70 synthesis, transcript stability or protein turnover. We therefore interpret recovery here operationally, for the individual markers and time points examined, rather than as complete physiological restoration.

The two cell lines provide a useful comparison because Kc cells express lamin Dm and lamin C, whereas S2 cells contain little or no detectable lamin C protein in the assay conditions described. Their different developmental origins and gene-expression programs provide additional sources of variation [9, 44, 45, 3]. The differences in Hsp70 kinetics and in some Top2, HSF and lamin Dm measurements are compatible with cell-context-dependent HS responses. However, these observations cannot attribute the differences specifically to lamin C without its targeted depletion or re-expression. RT-qPCR reference transcripts were selected after evaluation of stability under the tested conditions (Supplementary Figure 2); this is important because HS can affect commonly used housekeeping genes. The lack of statistically significant changes in several lamin-, Top2-and HSF-coding transcripts should likewise not be equated with an absence of post-transcriptional regulation.

### Lamin Dm forms, S25 phosphorylation and possible functional consequences

Earlier biochemical work established that Drosophila lamin Dm occurs as electrophoretically distinct forms, notably Dm0, Dm1 and Dm2, with more soluble mitotic or meiotic derivatives. These forms are linked to phosphorylation and changes in polymerization state [10, 11, 12, 13, 3]. In our Kc-cell samples, HS altered the relative intensity of the Dm1-and Dm2-associated bands and reduced binding of the monoclonal antibody ADL84, whose recognition is sensitive to phosphorylation within the S25-containing N-terminal epitope.

Crucially, Stuurman, Maus and Fisher [34] demonstrated restoration of ADL84 recognition after phosphatase treatment of lamin-containing blots, thereby providing direct support for the phosphorylation sensitivity of the antibody. The reproducible HS-associated decrease in ADL84 reactivity is consequently consistent with increased phosphorylation at S25; the response during recovery supports its transient nature.

The quantitative scope of this conclusion nevertheless needs to be defined carefully. An approximately 20% reduction in the ADL84-reactive signal, when confirmed by normalized densitometry, refers to an antibody-defined fraction relative to the corresponding control. It is not a direct measurement of the absolute percentage of all lamin Dm molecules carrying a phosphate at S25. Pre-existing phosphorylation differences may affect immunoblot quantification and epitope availability may affect the readout from immunofluorescence. There are also several technical issues possibly affecting precise readout of immunoblots: quality of electrophoretic separation of lamin Dm bands, material overloading, and the starting ratio of lamin Dm forms before HS may affect densitometric analyses. The calculated 20% of lamin Dm molecules change in S25 phosphorylation may appear a small and limited fraction of total lamin Dm. We need to take into consideration that most lamin molecules exist as polymerized filaments and are also involved in interactions with their nuclear partners; thus, the S25 epitope may be sterically less accessible for phosphorylation. The 20% of lamin Dm molecules additionally phosphorylated at S25 in HS theoretically may contribute significantly to the role that lamin plays during HS. The stoichiometric methods such as LC-MS/MS were frequently used for lamin Dm phosphosite mapping in Drosophila. Stoichiometric methods such as LC-MS/MS have frequently been used for lamin Dm phosphosite mapping in Drosophila [12, 15] but have yielded limited quantitative data; therefore, an epitope-specific antibody may be preferable when the demonstration of qualitative changes is sufficient.

The functional implications of S25 phosphorylation should also be assessed in light of earlier mutational studies. Phosphomimetic substitutions at lamin Dm S25, S45, T435 and S595 were associated with site-dependent changes in lamin properties, including polymerization, solubility or intracellular distribution [16]. Importantly, that study did not identify a general reduction of lamin Dm chromatin binding caused by individual phosphomimetic substitutions including S45 and T435 (N-and C-terminal “mitotic sites”). It would therefore be incorrect to use it as direct evidence that S25 phosphorylation decreases or increases chromatin affinity in vivo. The current data are more conservatively compatible with a change in the biochemical state or interaction environment of a subset of lamin Dm molecules, which may occur without a detectable change in bulk lamin localization.

S25 lies in the lamin Dm head region, within an R22PPS motif discussed as a potential site for 14-3-3-associated regulation; possible participation of prolyl isomerization by Pin1 and the influence of N-terminal phosphorylation on access to neighboring sites such as S45 are mechanistic hypotheses rather than demonstrated consequences of HS [13, 3]. Studies of mammalian lamin A/C have independently shown that N-terminal phosphorylation, including S22 phosphorylation during HS, can accompany stress-dependent nuclear adaptation in human cells and Drosophila respectively [46, 47]. This provides a conceptual parallel, but lamin A and Drosophila B-type lamin Dm are not interchangeable, and our immunofluorescence data did not demonstrate HS-induced redistribution of total lamin Dm. Thus, we propose phosphorylation-dependent changes in the properties or partners of a limited (at least 20%) lamin Dm population as a testable possibility, not as evidence of large-scale lamin disassembly or nuclear translocation.

### Top2 redistribution, chromatin organization and nuclear granules

A second important observation is that the HS-associated Top2 phenotype depends on the experimental system. In cultured Kc and S2 cells, diffuse nucleoplasmic Top2 staining was partly replaced by conspicuous Top2-positive granules (some associated with DAPI-negative regions) and heterogeneous nuclear staining (Figures 1A,B and 3A). In stage 10–12 embryonic nuclei, by contrast, the quantified effect was Top2 enrichment near the nuclear periphery (Figure 3B–E). Peripheral accumulation of DAPI-stained chromatin was demonstrated separately in polyploid larval nuclei (Figure 3F,G). Because the larval experiment did not establish accompanying peripheral Top2 relocation in the same nuclei, these observations cannot demonstrate coordinated movement of Top2-bound chromatin to the lamina. They nevertheless identify distinct, potentially related aspects of nuclear adaptation to HS and justify future experiments using simultaneous Top2, lamin and DNA-domain measurements in the same cells, interactome, ChIP-seq, and RNA-seq experiments.

Other immunofluorescence observations support the existence of a broader stress response but cannot determine the mechanism of Top2 redistribution. Hsp90 displayed the expected shift toward nuclear staining in Kc cells during HS (Supplementary Figure 3). The changes reported for HSF and other nuclear antigens may reflect stress-altered signaling. The absence of a clear change in total lamin Dm staining is especially relevant: Top2 can change its nuclear pattern without an accompanying microscopically detectable redistribution of the bulk lamin polymer.

The heterogeneous behavior of Top2 is consistent with earlier work by Fisher and colleagues. Different Drosophila Top2 fractions were characterized by their extraction properties, phosphorylation, catalytic activity and associations with DNA or RNA. In particular, the P-10 and P-130 biochemical populations are not equivalent; the more highly phosphorylated fraction displayed reduced catalytic activity and distinct nucleic-acid associations, while a non-chromosomal population was especially associated with RNA [18, 21, 19, 20, 22]. The C-terminal region of Top2 contributes to its nucleic-acid associations and phosphorylation-dependent behavior [22]. It is therefore plausible that a restricted pre-existing Top2 subpopulation, rather than the entire cellular Top2 pool, changes distribution or partners after HS.

There is also independent evidence that HS changes Top2 activity or occupancy at particular chromatin loci. Rowe, Wang and Liu [23] mapped HS-responsive changes in Top2 cleavage sites, and Udvardy and Schedl [24] described redistribution of Top2 sites of action at the 87A7 heat-shock locus during induction and recovery. These results provide a biologically relevant precedent for dynamic Top2–chromatin associations and help explain why the present nuclear redistribution merits investigation. They do not establish a global increase in Top2 binding or show that the Top2-positive granules observed here are heat-shock loci. The possibility that some Top2 granules are RNA-rich nuclear bodies is suggested by earlier observations of RNA-associated Top2 fractions, but the granules have not been characterized for RNA, RNP proteins or established stress-body markers. Stress-responsive RNP granules do occur in Drosophila (cytoplasmic only reported so far) and other invertebrates [48]; nevertheless, these particular Top2-positive structures should not be designated stress granules without direct molecular evidence [49, 50].

### Salt extractability as an indicator of biochemical remodeling

Earlier studies described HS-dependent enrichment of lamins, Top2 and other nuclear proteins in extraction-resistant or karyoskeletal preparations [51, 52, 25, 26, 19, 22]. Such findings were historically interpreted through the nuclear-matrix concept. However, an insoluble fraction is operationally defined by the extraction protocol and cannot, by itself, prove that a corresponding structural scaffold exists in intact cells. Salt resistance may be influenced by polymerization, multiprotein and nucleic-acid interactions, post-translational modifications and, under stress, aggregation. For this reason, the current measurements are described as changes in extractability, without treating the 500 mM NaCl fraction as a direct image of a physiological nuclear matrix.

Sequential extraction using 50, 150 and 500 mM NaCl provided a comparative biochemical view of HS responses in Kc and S2 cells (Figure 4; Supplementary Figure 5). These concentrations recover overlapping sets of proteins rather than sharply separated biological compartments. Moreover, our densitometric comparisons were performed within, not across, each fraction. The data can therefore identify differences between N and HS samples extracted under a particular condition but cannot independently calculate the redistribution of an entire protein population between all fractions. This limitation is important when linking these data to microscopy.

Despite these restrictions, the extractability results provide a second line of evidence that HS affects the molecular context of lamin Dm and Top2. In Kc cells, lamin Dm recovery increased in the intermediate-and high-salt fractions, whereas Top2 declined in the 50 mM fraction and increased in the 150 mM fraction; HDAC1 and HSF showed additional shifts (Figure 4A). In S2 cells, some responses fully overlapped with those in Kc cells, especially for HSF and HDAC1, while the affected lamin Dm and Top2 fractions partly overlapped (Figure 4B). None of the tested fractions changed in opposite direction which indicates strong similarity of proteins response in HS in both cell lines and common mechanisms modifying their properties. This correlated with similar relocation of Top2 and S25 phosphorylation. Altered ADL84 reactivity in a Kc low-salt fraction is consistent with heterogeneity of the S25-sensitive lamin population. Increase in solubility of HSF and HDAC1 in both cell lines are in agreement with their known response to HS. Taken together, the changes are compatible with modulation of protein assembly, nucleic-acid association or intermolecular contacts during HS, not necessarily with movement of the same complexes between defined nuclear compartments. An alternative explanation might be potential specific-protein-fraction heat-induced aggregation or extraction-induced loss/increase of solubility. However unlikely that might be, such possibility cannot be excluded.

### Nucleic-acid association of lamin Dm and Top2: evidence and assay limitations

The in vivo photocrosslinking results provide another mechanistically relevant dimension because both proteins were previously shown to associate with nucleic acids. Lamin Dm binds detectable DNA and RNA fractions in interphase, but lamin C did not yield comparable signals in the same experimental system [17]. Top2 binds DNA and RNA in different cell-cycle and biochemical populations, including the RNA-associated forms described by Rzepecki and Fisher [22]. There is no doubt that lamin Dm (B-type lamin) binds DNA as all B1, B2-type lamins do. Lamin C do not bind DNA as all lamin C do. Protein-DNA or specifically Top2-DNA crosslinking were used for protein-DNA complexes maping, also in vivo. Our BrdU/UV photocrosslinking and immunoprecipitation experiments therefore build on established assays rather than assigning a previously unknown capacity for nucleic-acid binding. The observed increase in radiolabeled, protein-associated material after HS is consistent with enhanced recovery of a nucleic-acid-associated subset of lamin Dm and Top2 (Figure 6C–E). Because only limited fractions of the total protein populations contribute to the signal, probably located in close proximity, however, the result cannot be generalized to every lamin Dm or Top2 molecule.

The chemistry of Top2 introduces an additional interpretative consideration. In its catalytic cycle, Top2 transiently forms a covalent protein–DNA cleavage intermediate; SDS-based lysis can preserve such complexes, and catalytic Top2 cleavage sites have long been exploited to map the enzyme’s action in Drosophila chromatin [23, 24, 53, 54]. Consequently, the Top2-associated DNA recovered here may include both UV-crosslinked contacts and covalent catalytic intermediates. These contributions cannot be separated using the current assay and the yield of contribution of each is irrelevant for the experiment. Potential risk of stress induction associated with UV exposure, DNA damage-associated relocalization, DNA breaks-associate chromatin relocalization of heterochromatin and genotoxic stress induction (Ryu et al. 2015,Nature Cell Biology;, 17(11), 1401–1411) can be excluded as an important contributory. Report demonstrates that Kc irradiated with 5G dose X-ray generated 12-18 double strand breaks in heterochromatin (inside cell nuclei) associating homologous recombination repair complex components including TopBP1. Statistically only up to 1 foci per nucleus of such complexes relocates to nuclear periphery upon irradiation. Considering recruitment of Top2 to such complex, the potential contribution to the Top2 – lamin Dm chromatin complexes increase in co-IP, PLA and crosslinking analyses such mechanism may contribute only a minor addition at the most to our demonstrated HS induction. Besides, used irradiation method creates many chromosomal aberrations and does not contribute to the demonstrated lamin Dm-Top2 complexes in N conditions. No Hsp70 induction by our procedure eliminates the question of BrdU feeding and crosslinking procedure as a potential stress activation under N conditions. Additionaly, nuclease sensitive co-IP of lamin Dm and Top2 was demonstrated in N conditions, HS only increased the co-IP and co-localization in PLA assay.

The two sets of nuclease experiments should be regarded as complementary rather than contradictory. Under native extraction conditions, lamin Dm and Top2 co-immunoprecipitated reciprocally, and additional on-bead S7 treatment did not visibly eliminate their association (Figure 5D,E). This is compatible with complexes that retain protein-mediated contacts or are protected from nuclease digestion. After crosslinking and the different extraction (denaturing conditions) or purification steps used for Figure 6A,B, S7 treatment reduced Top2 recovery with lamin Dm and diminished the radiolabeled material. This indicates a nuclease-sensitive component under those conditions. A reasonable working interpretation is that the assays recover overlapping but not identical populations: some associations may be largely nucleic-acid-independent, whereas others require accessible nucleic-acid fragments or are stabilized by them. Neither experiment alone establishes direct lamin Dm–Top2 binding, and differences in nuclease accessibility, immobilization, crosslinking and washing must be considered before estimating the relative sizes of these populations. Nevertheless, the demonstrated Top2-lamin Dm reciprocal co-IP under native, paraformaldehyde crosslinking and BrdU-UV crosslinking conditions and their increase in HS was established beyond doubt.

Similarly, decreasing radiolabel with increasing nuclease treatment (Figure 6B) is expected for protein-associated nucleic acids. The reduction of lamin Dm signal during prolonged on-bead digestion might reflect loss of complexes retained through nucleic acids or a change in recovery during washing. It cannot be assigned uniquely to release of a particular lamin Dm–Top2–DNA bridge. Weak radioactivity near protein-rich or IgG bands also requires cautious interpretation because residual nucleic-acid fragments and labeling background can contribute to autoradiographic signals. These are reasons to emphasize nuclease-sensitive, immunoprecipitated material rather than to infer a precise molecular topology from band positions.

DNase I and RNase A treatment produced broadly similar relative sensitivity patterns in control and HS samples (Figure 6C). This is consistent with contributions from both classes of nucleic acid but does not demonstrate that the exact DNA sequences, RNA species or binding modes remained unchanged. The 150 mM NaCl-soluble material gave prominent nucleic-acid-associated signals in the fractionation analysis (Figure 6D), reinforcing the existence of a biochemically distinct crosslinkable pool. Most importantly, comparisons of N, HS and recovery conditions in Figure 6E showed increased nucleic-acid-associated signals after HS and a subsequent decline. This supports a dynamic HS response for a limited pool of lamin Dm and Top2 but is not sufficient to conclude that an unchanged protein complex has moved bodily from the nuclear interior to the lamina.

The distamycin experiment offers an additional, although indirect, mechanistic argument. Minor-groove-binding compounds have been used previously to investigate the interaction of lamin B1 with DNA, including matrix-attachment-region substrates [55]. In our Kc-cell assay, distamycin decreased the lamin Dm-associated signal under both normal and HS conditions and modified the Top2-associated signal more prominently after HS (Figure 6F). The findings suggest that the recovered complexes are sensitive to changes in the accessibility or conformation of DNA regions affected by the drug. They do not establish direct minor-groove recognition by either protein, identify AT-rich binding sequences or prove that HS alters DNA sequence specificity. Competitive occupancy of DNA, altered chromatin accessibility, and changes in UV-crosslinking efficiency are alternative explanations. A follow-up experiment with ChIP-seq for Top2 and lamin Dm would solve the sequence mapping and LAD binding. Preliminary experimental data confirm HS-mediated Top2 relocation on chromosomes.

The partial agreement between Kc and S2 responses is also informative. These lines differ in lamin C expression [9], developmental history and basal gene-expression programs, yet they show HS-dependent changes affecting related classes of nuclear proteins. This is compatible with a conserved component of the general stress response superimposed on cell-specific regulatory networks [56, 2, 3]. The cell-specific magnitudes and affected extraction fractions caution against assuming that the same lamin Dm or Top2 species responds identically in both lines. Phosphorylation measurements should be interpreted separately for the relevant antibody, cell type and fraction; differences between whole-cell and salt-extracted samples are not necessarily inconsistent, because the populations examined are not the same.

### Lamin Dm–Top2 proximity, co-immunoprecipitation and the meaning of a restricted interacting pool

PLA signals, native and PFA crosslinking co-IP and the nucleic-acid assays converge on an important but bounded conclusion: lamin Dm and Top2 can occur in shared molecular environments. PLA revealed proximity signals in Kc cells and third-instar larval material, including many foci in polytene salivary-gland nuclei and fewer foci in diploid cells (Figure 5B,C,H,I). A low number of foci per diploid nucleus does not refute a biologically relevant association. The two proteins are abundant in largely distinct compartments, the assay samples only molecules within its detection distance and epitope-accessibility constraints, and the number of chromosomal copies differs dramatically between polytene and diploid nuclei. Polytene foci therefore cannot be extrapolated quantitatively to Kc or imaginal-disc cells; equally, a restricted diploid-cell interaction need not be ubiquitous around the entire nuclear rim. These possibilities account for the qualitative pattern but should be tested through properly normalized foci-per-nucleus distributions and matched nuclear-volume or DNA-content analyses.

Reciprocal co-immunoprecipitation supports the presence of lamin Dm–Top2-containing complexes in native extracts (Figure 5D,E), and PFA-assisted extraction permits recovery of additional crosslinked material (Figure 5F,G). Nevertheless, the present Figure 5F cannot independently establish a statistically significant HS-induced increase in direct protein–protein binding. Increased signal could arise from differences in extraction, crosslinking, epitope accessibility or the abundance of recovered complexes. PLA foci increase during HS, and Figure 6E shows increased nucleic-acid-associated signal; these measurements are consistent with a change in proximity or in the fraction of molecular assemblies that can be detected, rather than proof of a stronger direct lamin Dm–Top2 interaction. Top2 relocation toward the nuclear periphery in cells and embryos favors the increased proximity hypothesis. The comparable lamin Dm band pattern in some Top2 co-immunoprecipitates might suggest that both electrophoretic forms are recovered, but it does not establish that S25 phosphorylation has no effect on their molecular affinity.

### A testable working model and the limits of the present evidence

Taken together, the phosphorylation-sensitive lamin Dm changes, Top2 nuclear phenotypes, salt-extractability shifts, PLA proximity and increased nucleic-acid-associated signals support a working model of HS-dependent remodeling of limited nuclear protein populations. Rather than assuming one concerted translocation event, the model allows several, not mutually exclusive possibilities: altered phosphorylation and assembly of a lamin Dm subset, changes in the partner proteins or nucleic acids associated with Top2, and increased opportunities for the two proteins to approach one another within some nuclear complexes. This interpretation is biologically plausible because Top2 has a heterogeneous biochemical repertoire and lamin Dm participates in multiprotein networks, but its specific molecular sequence remains to be established.

Known links among lamins, LEM-domain proteins, BAF, HP1, LBR, nuclear-pore proteins and chromatin provide a structural context in which such remodeling could occur [42, 57, 58, 3]. Nucleoporins, including Nup98, Nup107 and Elys, have been implicated in nuclear architecture and gene-regulatory interactions, while broader studies of LADs and TADs demonstrate the importance of three-dimensional chromatin organization [35, 36, 44, 39]. These published relationships make it reasonable to ask whether HS alters contacts among Top2, lamin Dm and selected chromatin domains and domain regulators. They do not show that the particular Top2-positive structures reported here are associated with nucleoporins, constitutive LADs or new facultative LADs. Identifying such contacts would require locus-specific imaging, RNA-seq and ChIP-seq or other spatially resolved genomic methods.

An additional aspect worth retaining is the possible contribution of lamin composition to these interactions. S2 cells and early embryonic cells differ from Kc cells in the relative presence of lamin C, and lamin C can influence nuclear mechanics and the organization of lamin-associated complexes [59, 60, 3]. The in vivo nucleic-acid-binding difference between lamin Dm and lamin C observed by Rzepecki et al. [17] supplies a testable rationale for comparing these backgrounds. However, lamin C expression is not the only difference between the models, and the experiments do not demonstrate that absence of lamin C causes faster Hsp70 attenuation, Top2 granule formation or altered extractability. Manipulating lamin C in a matched genetic background would be necessary to distinguish a lamin-dependent effect from developmental or lineage-specific differences.

A more ambitious extension of the model would involve HS-dependent repositioning of particular chromatin loci toward the lamina, potentially with alterations in Top2-associated complexes. This proposal is consistent with known lamina–chromatin interactions and with reports of HS-dependent Top2 cleavage-site redistribution [23, 24], but it is not demonstrated by the present experiments. No specific transcriptionally repressed loci were mapped, no formation of facultative LADs was measured, and the Top2 and chromatin redistribution images were obtained from different experimental contexts. Similarly, transient retention of non-HS transcripts near nuclear pore complexes, or incorporation of Top2 into nuclear stress bodies, remains speculative and should not be presented as an explanation already supported by the figures. These ideas can motivate targeted analyses of gene loci, RNA species, and nuclear-pore or stress-body markers in future work.

The strengths of this study lie in the convergence of imaging, phospho-sensitive immunodetection, biochemical fractionation, PLA and in vivo nucleic-acid assays, each addressing a different part of the response. In conclusion, the data support transient, model-dependent changes in the properties and molecular neighborhoods of lamin Dm and Top2, together with enhanced nucleic-acid association of restricted protein fractions. They do not yet establish that a particular, direct lamin Dm–Top2 interaction increases during HS or that specific genes are relocated and transcriptionally regulated through the proposed lamina-associated mechanism.

Supplementary video 1

### PLA movie1

The Proximity Ligation Assay PLA for lamin Dm and Top2 in polytenic nuclei of the 3rd instar larvae salivary gland cells. Colocalization signal visible as re loci. Proteins were labeled in PLA via the anti-lamin Dm monoclonal ADL67 antibody and affinity-purified anti-Top2 antibodies.

Supplementary video 2

### PLA movie2

The Proximity Ligation Assay PLA for lamin Dm and Top2 in polytenic nuclei of the 3rd instar larvae salivary gland cells. Colocalization signal visible as re loci. Proteins were labeled in PLA via the anti-lamin Dm monoclonal ADL67 antibody and affinity-purified anti-Top2 antibodies.

## Author Contributions

The authors confirm the contributions to the paper as follows: funding support, project administration (PI), concept of the study, general supervision, original draft and manuscript writing: RR; work supervision, direct supervision and experiment planning: RR, MR, AT, JJ, MM, KP, AZ; conducting research, experimental work, data analysis and visualization: MR, AT, JJ, AZ, MM, RR. Manuscript editing and figure preparation RR, MR, AT, JJ, AZ, MM. All the authors reviewed the results and approved the final version of the manuscript.

## Funding

Funded by the National Centre for Science (NCN) grant No. 2016/21/B/NZ4/00541.

## Data Availability Statement

Data associated with this manuscript have been deposited on a server at the Faculty of Biotechnology, University of Wroclaw. Please contact the authors for information on access to the data.

## Acknowledgments

I would like to thank Prof. Paul A. Fisher (Dept. of Pharmacology, SUNY SB, USA) for allowing me to perform initial studies in his laboratory.

## Conflicts of Interest

The authors declare that they have no competing interests.

## Supplementary Materials

**Supplementary Figure 1.**
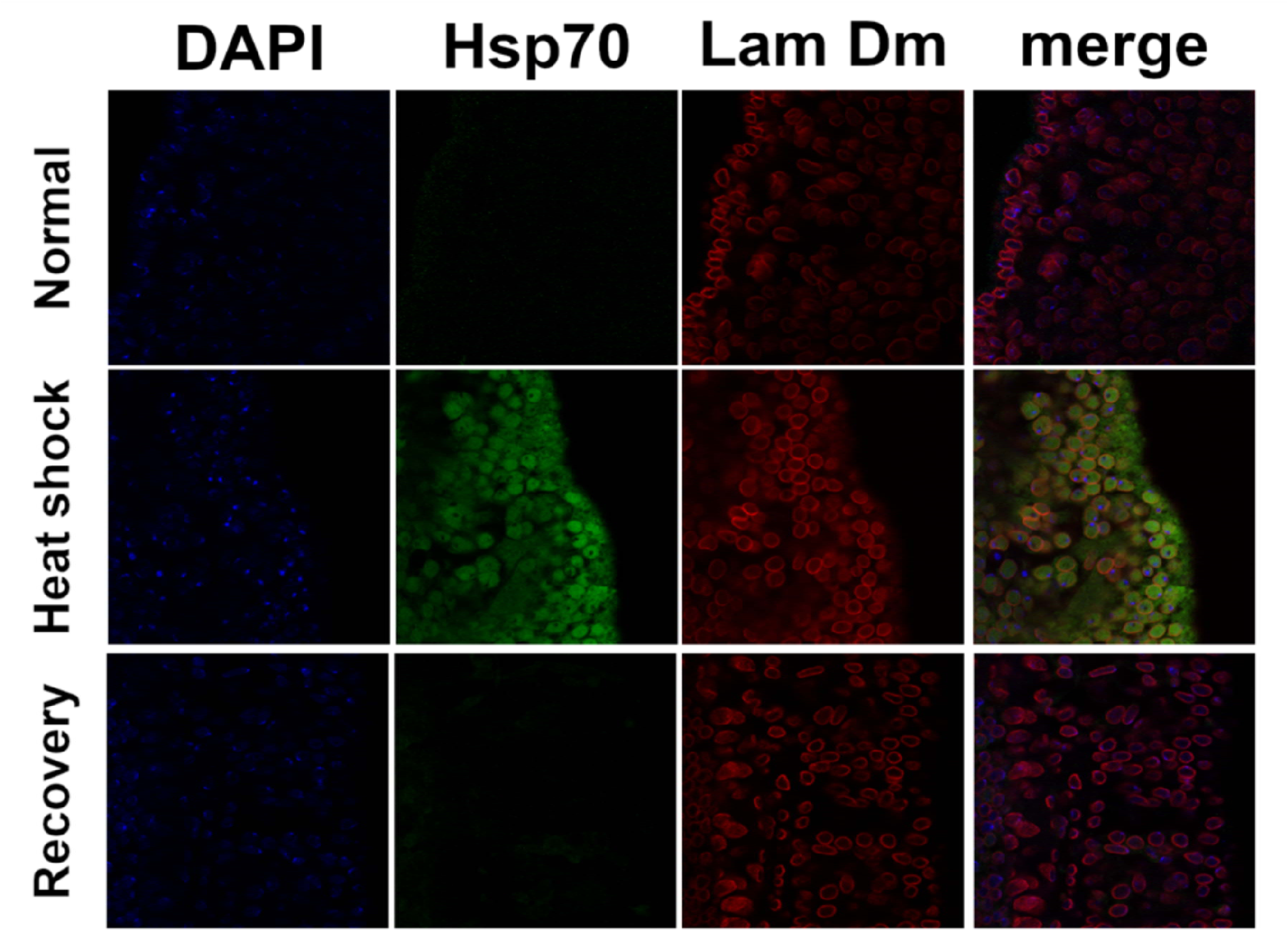
Heat shock induction in *D. melanogaster* embryos. The efficiency of heat shock induction and recovery was examined via immunofluorescence staining of *D. melanogaster* embryos under different conditions: normal, heat shock (1 h) and recovery. Merge, chromatin (DAPI - blue), lamin Dm (Lam Dm - red), and heat shock induction (Hsp70 - green) visualization.

**Supplementary Figure 2.**
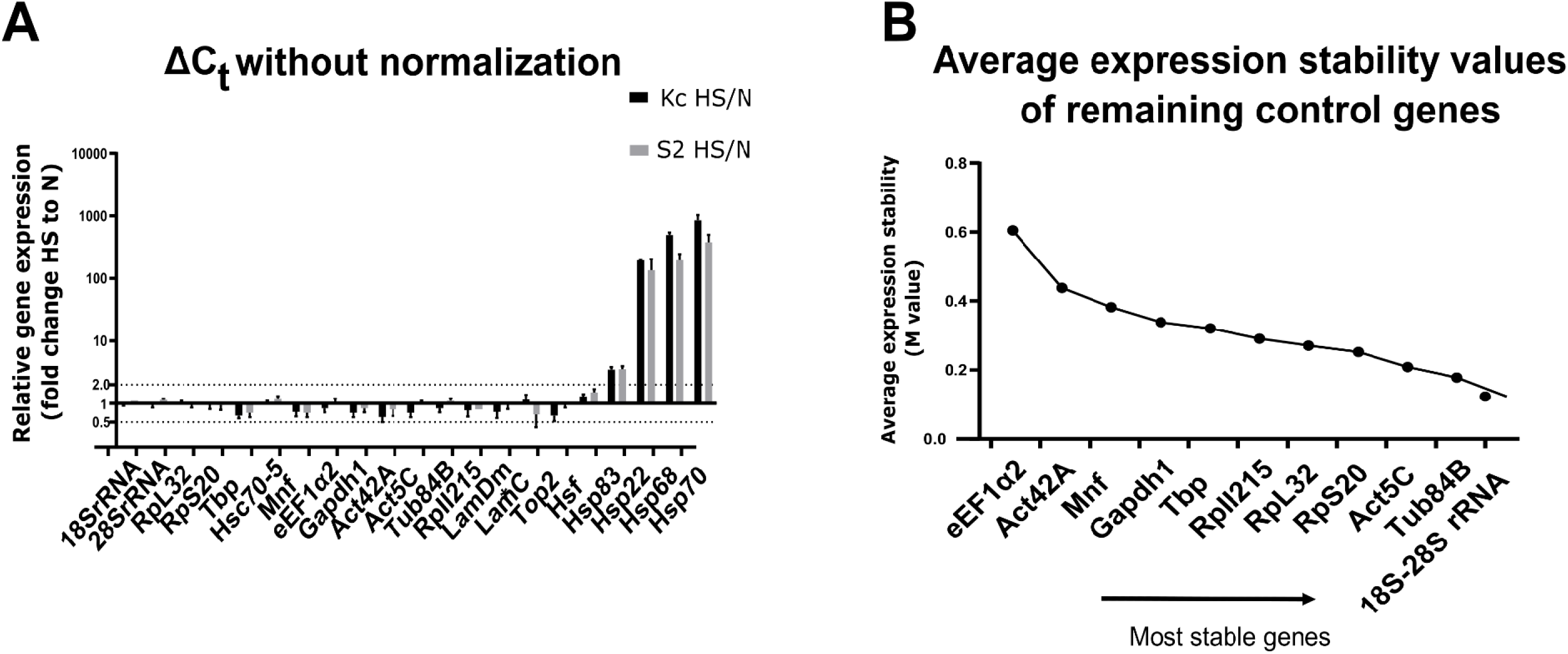
Assessment of gene stability after heat shock induction in *Drosophila melanogaster* cells (Kc and S2) after 1 h of HS. Gene expression of all tested genes (names given on the x-axis) in Kc and S2 cells (A). Bars represent the fold change in the transcript level after 1 h of heat shock induction for Kc (black) and S2 (gray). Only normalization to the control (N) was used without further normalization to the housekeeping genes. The threshold marked by a dotted line was set as 2-fold. Determination of the most stable reference (housekeeping) genes from a set of tested candidate reference genes was performed via the geNorm algorithm (B). The average expression stability was measured to choose the common endogenous control for the Kc and S2 cell lines upon heat shock.

**Supplementary Figure 3.**
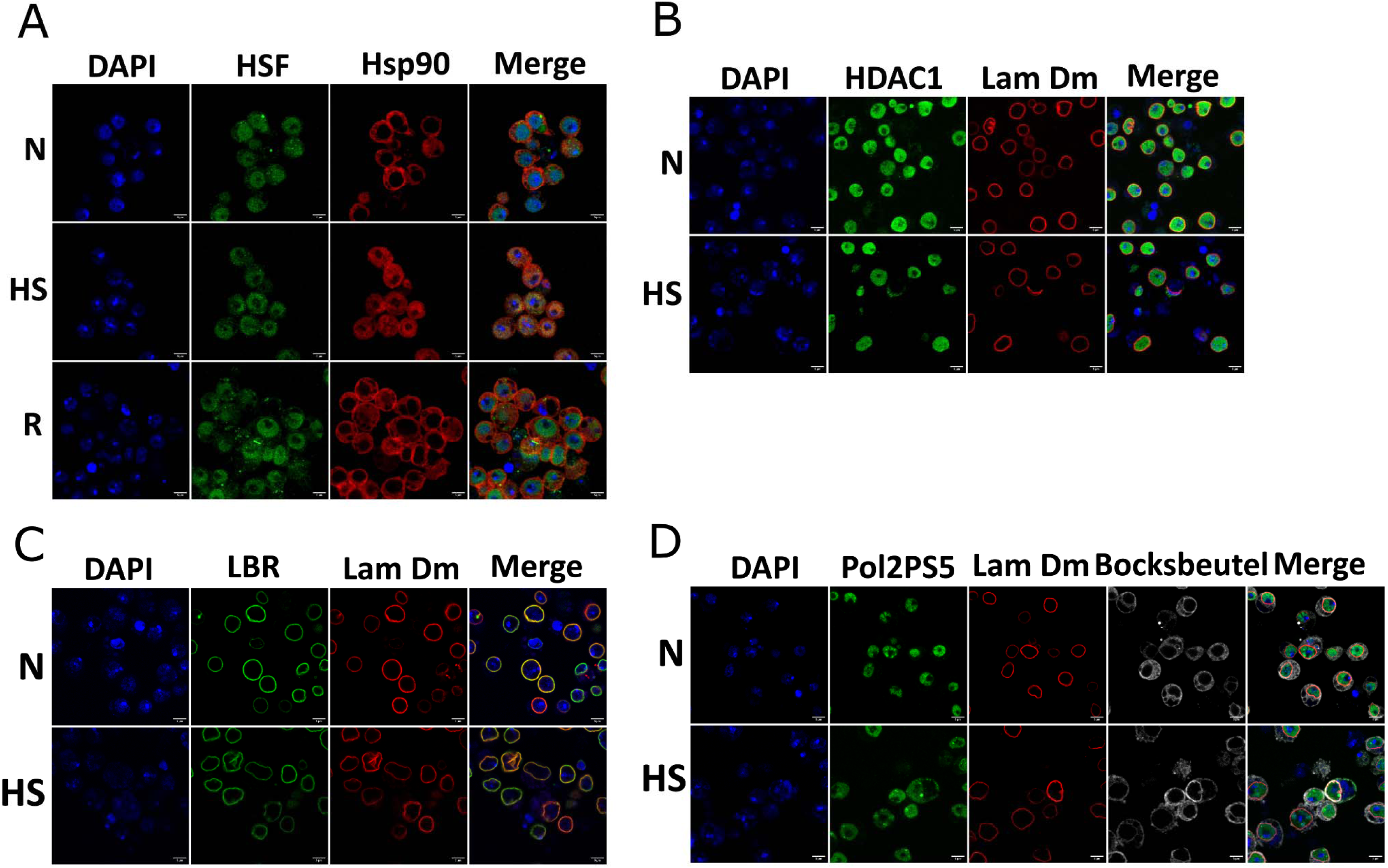
Immunofluorescence analysis of the distribution of nuclear proteins in Kc cells. Immunofluorescence staining in Kc cells; control cells (N), heat shock induced cells (HS) and recovery (R). Immunofluorescence staining for HSF, HDAC1, LBR and RNA Polymerase 2 pS5 (all green) and Hsp90 and lamin Dm (all red). DNA stained with DAPI (blue). Please note the expected, transient relocation of Hsp90 protein to the cell nuclei of Kc cells in HS and concomitant relocation of HSF protein to the cell nuclei in HS. No other significant changes have been detected. Scale bars = 5 µm.

**Supplementary Figure 4.**
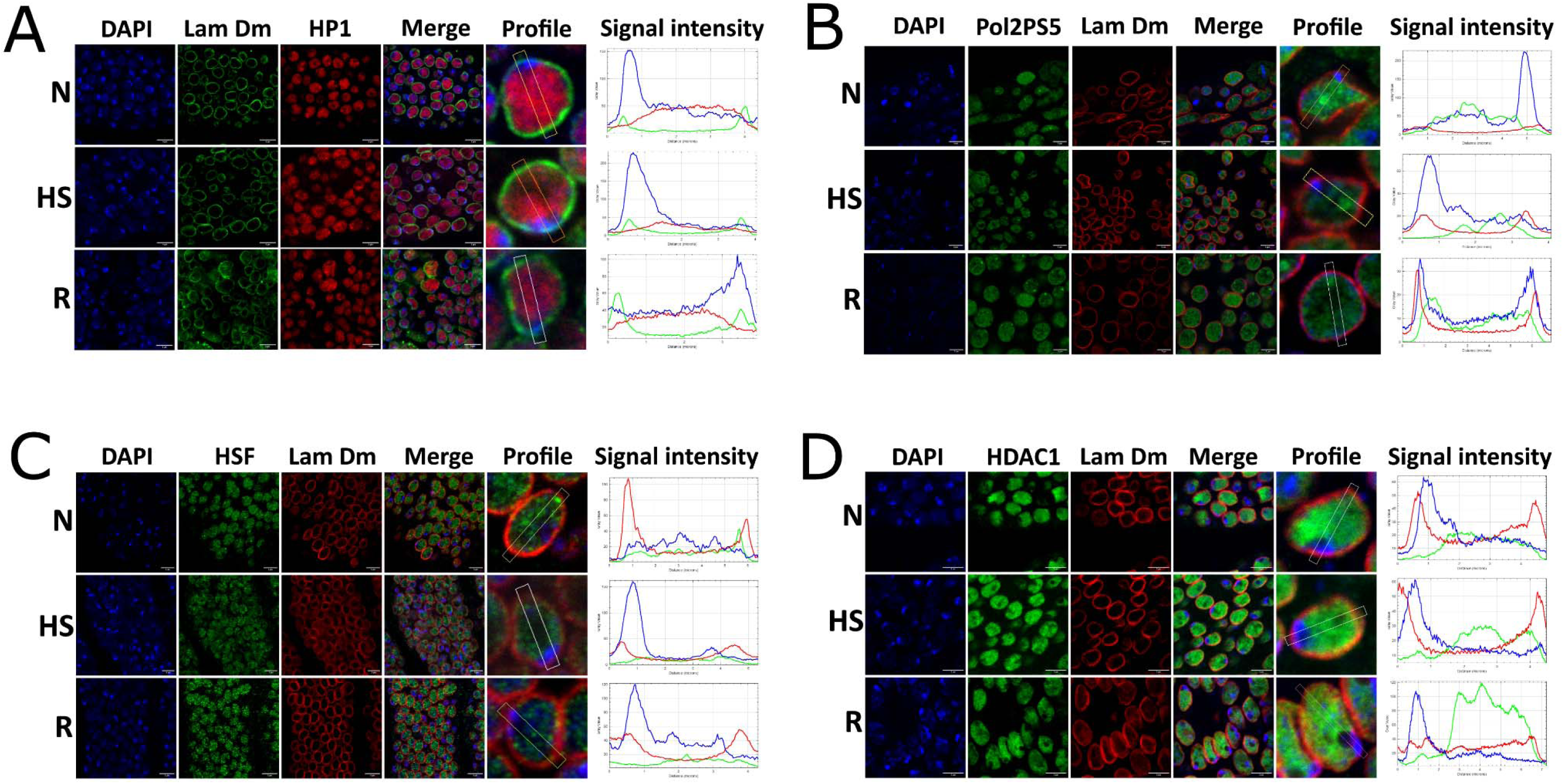
Immunofluorescence analysis of the distribution of other nuclear proteins in D. melanogaster embryos. Immunofluorescence staining of *D. melanogaster* embryos under N, HS (1 h) and R conditions with additional fluorescence intensity analysis (A-D). Distribution of HP1 (A – red), Pol2PS5 (B - green), HSF (C - green) and HDAC1 (D – green; DNA was stained with DAPI (blue); lamin Dm (A)–green, (B-D)-red. was visualized. The area from which the fluorescence intensity was analyzed is marked with a yellow line in the “Profile” column on each graph. A graph of the fluorescence intensity distribution (Signal intensity) for each channel is also presented (for each staining separately in the corresponding color). No significant redistribution was observed under the examined conditions. Scale bars = 5 µm.

**Supplementary Figure 5.**
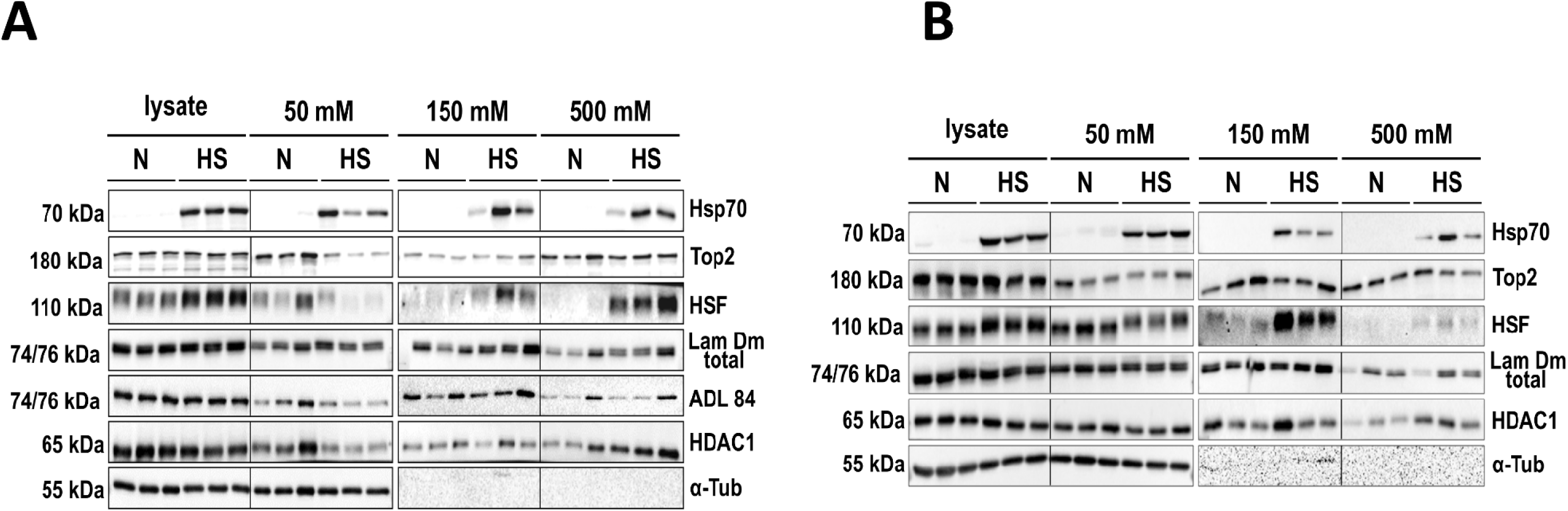
Western blot analysis of the solubility of proteins in Kc and S2 control cells and after heat shock induction. Typical western blot from experiments for Figure 4 was shown. Panel A-Kc cells; Panel B-S2 cells. The experiment was conducted in the same way as in Figure 4. The solubility tests were performed for Hsp70, lamin Dm (total lamin Dm fraction AP rabbit Ab anti lamin Dm), lamin Dm unphosphorylated on S25 (with ADL84), Top2, HDAC1, HSF and αTub as a loading/normalization control. Please note nicely visible both forms of lamin Dm: Dm1 and Dm2 in extracts and less visible/resolved in lysate fractions and nice correlation of increase of upper, lamin Dm2 form in extracts from heat shocked Kc and less pronounced in S2 cells (A and B). Please note the increase of lamin Dm single band staining with ADL84 antibodies in control extract (50 mM NaCl) from control Kc cells indicated increased level of lamin Dm1 form in this extract (as in Figure 4A (ADL84).

---

**Supplementary Table 1.**
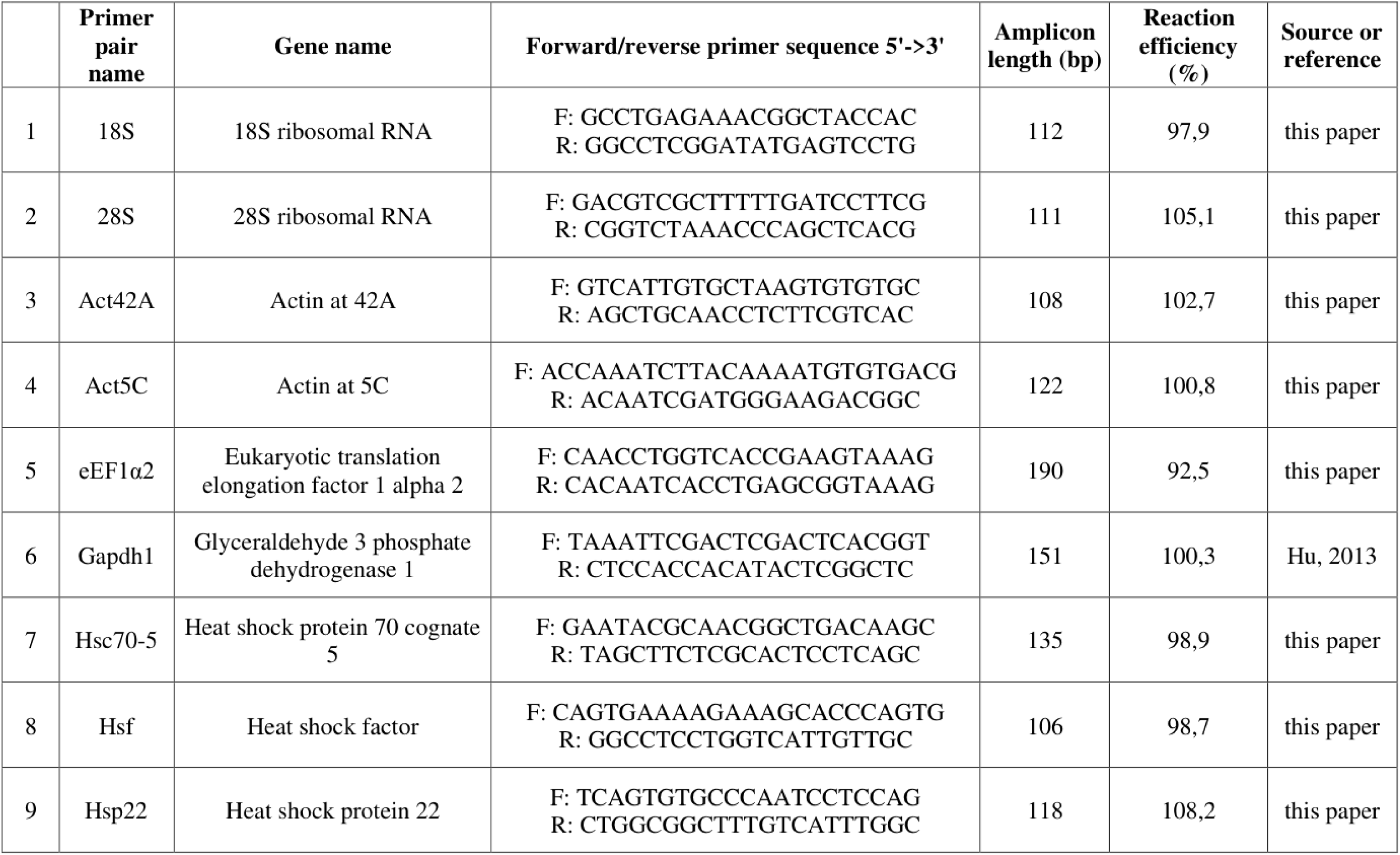
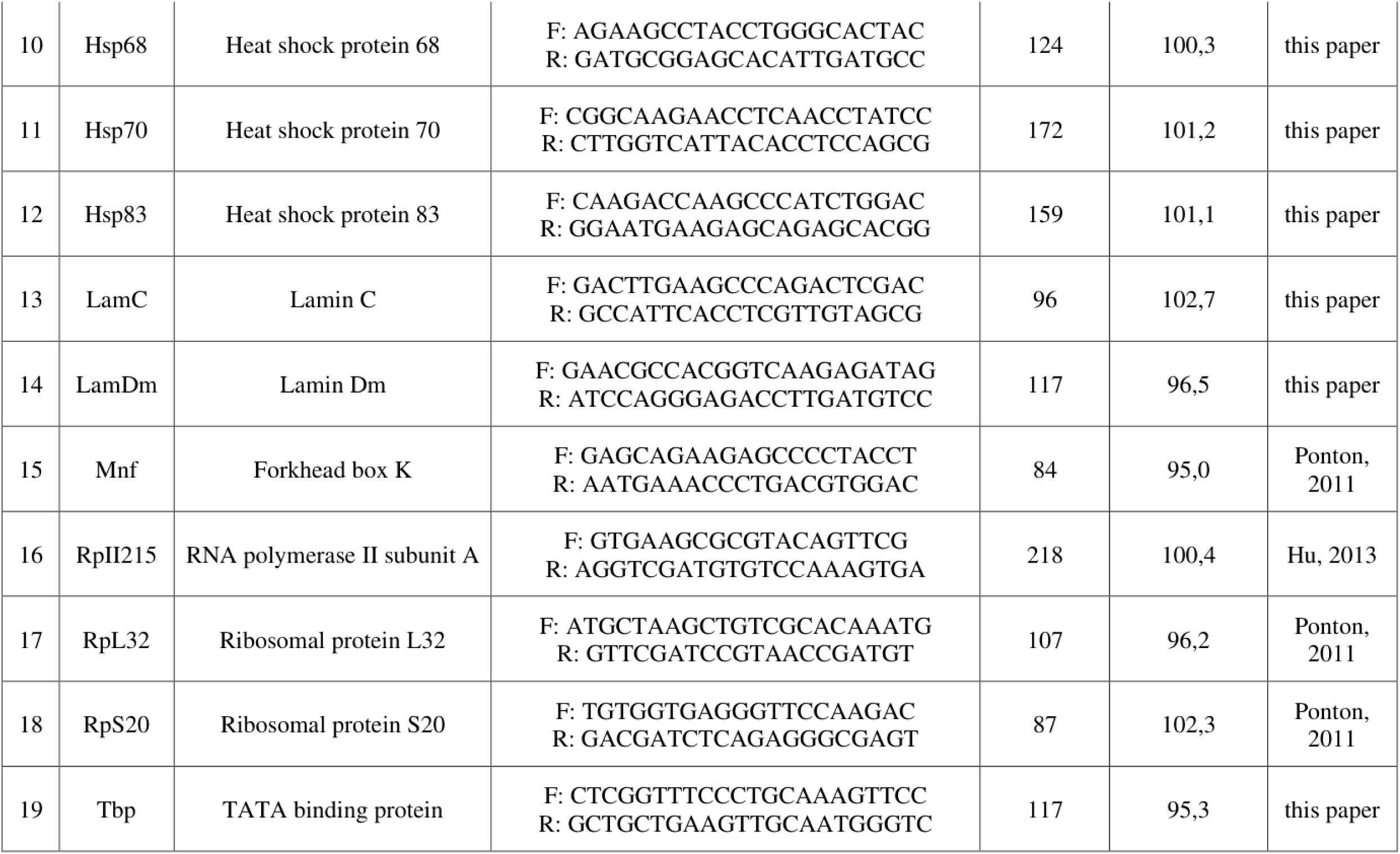
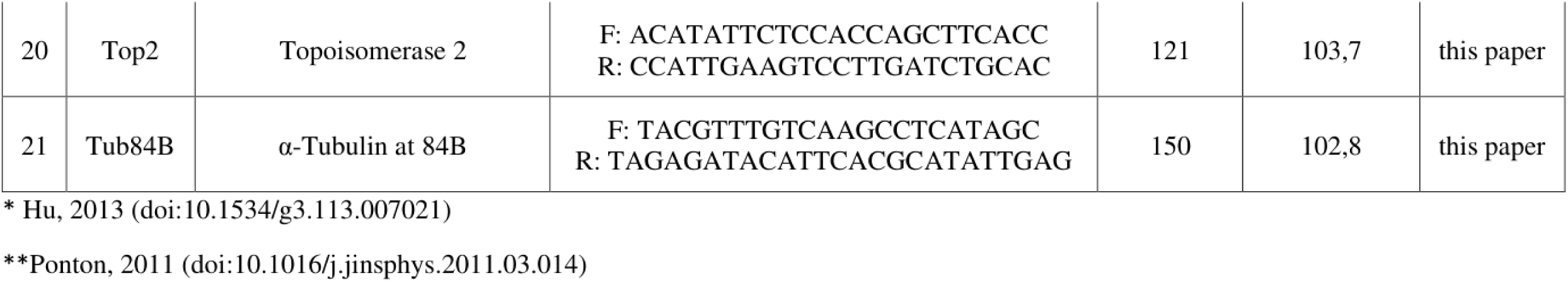
List of the primers used for RT qPCR analysis. Table with a list of all primers used in the RT qPCR analyses. Their sequences, efficiency and source.

## Notes

### Competing Interest Statement

The authors have declared no competing interest.

### Summary of Updates

Abstract thesis Introduction content and layout, shortening Revised Results section, new layout and description, revised thesis and documentation Major revision of Discussion section Revised and updated Reference list

